# Structural basis of a two-antibody cocktail exhibiting highly potent and broadly neutralizing activities against SARS-CoV-2 variants including diverse Omicron sublineages

**DOI:** 10.1101/2022.05.27.493682

**Authors:** Xiaoman Li, Yongbing Pan, Qiangling Yin, Zejun Wang, Sisi Shan, Laixing Zhang, Jinfang Yu, Yuanyuan Qu, Lina Sun, Fang Gui, Jia Lu, Zhaofei Jing, Wei Wu, Tao Huang, Xuanling Shi, Jiandong Li, Xinguo Li, Dexin Li, Shiwen Wang, Maojun Yang, Linqi Zhang, Kai Duan, Mifang Liang, Xiaoming Yang, Xinquan Wang

**Affiliations:** The Ministry of Education Key Laboratory of Protein Science, Beijing Advanced Innovation Center for Structural Biology, Beijing Frontier Research Center for Biological Structure, School of Life Sciences, Tsinghua University, Beijing 100084, China; National Engineering Technology Research Center for Combined Vaccines, Wuhan Institute of Biological Products Co. Ltd., Wuhan 430070, China; State Key Laboratory for Molecular Virology and Genetic Engineering, National Institute for Viral Disease Control and Prevention, Chinese Center for Disease Control and Prevention, Beijing 102206, China; NexVac Research Center, Comprehensive AIDS Research Center, Center for Infectious Disease Research, Department of Basic Medical Sciences, School of Medicine, Tsinghua University, Beijing 100084, China; Institution of Infectious Diseases, Shenzhen Bay Laboratory, Shenzhen 518107, China; CDC-WIV Joint Research Center for Emerging Diseases and Biosafety, Wuhan 430071, China

## Abstract

SARS-CoV-2 variants of concern (VOCs), especially the latest Omicron, have exhibited severe antibody evasion. Broadly neutralizing antibodies with high potency against Omicron are urgently needed for understanding working mechanisms and developing therapeutic agents. In this study, we characterized previously reported F61, which was isolated from convalescent patients infected with prototype SARS-CoV-2, as a broadly neutralizing antibody against all VOCs including Omicron BA.1, BA.1.1, BA.2, BA.3 and BA.4 sublineages by utilizing antigen binding and cell infection assays. We also identified and characterized another broadly neutralizing antibody D2 with epitope distinct from that of F61. More importantly, we showed that a combination of F61 with D2 exhibited synergy in neutralization and protecting mice from SARS-CoV-2 Delta and Omicron BA.1 variants. Cryo-EM structures of the spike-F61 and spike-D2 binary complexes revealed the distinct epitopes of F61 and D2 at atomic level and the structural basis for neutralization. Cryo-EM structure of the Omicron-spike-F61-D2 ternary complex provides further structural insights into the synergy between F61 and D2. These results collectively indicated F61 and F61-D2 cocktail as promising therapeutic antibodies for combating SARS-CoV-2 variants including diverse Omicron sublineages.

## Introduction

Since the first documented cases of the SARS-CoV-2 infection in Wuhan, China in late 2019, the COVID-19 pandemic has been posing a severe threat to the global public health, with more than 523 million infections and over 6 million deaths around the world^1–3^ (https://www.who.int/emergencies/diseases/novel-coronavirus-2019/situation-reports/). Vaccines, monoclonal neutralizing antibodies, small-molecule drugs have been successfully developed for prophylaxis and treatment in fighting against the SARS-CoV-2^3–20^. However, SARS-CoV-2 variants, especially variants of concern (VOC) with changed pathogenicity, increased transmissibility and resistance to convalescent/vaccination sera and monoclonal antibodies have emerged repeatedly during the circulation^21–24^. In 2020, the first VOC Alpha (B.1.1.7) was identified in the United Kingdom^25,26^, followed by Beta (B.1.351) in South Africa^27^ and Gamma (P.1) in Brazil^28^. These three VOCs mainly circulated in their identified and neighboring countries. In contrast, Delta (B.1.617.2) first detected in India in late 2020 quickly spread to nearly all countries and became the global dominant VOC in 2021^29–32^. In November 2021, Omicron (B.1.1.529) was reported from South Africa, and the World Health Organization (WHO) immediately designated it as the fifth VOC due to its over 40 mutations in the spike (S) glycoprotein, at least three times more than the number found in previous four VOCs^33–36^. Although Omicron has lower fatality rate than Delta, it quickly outcompeted Delta and became the dominant circulating variant in 2022, due to the significantly increased transmissibility^33,37^. Currently, the major sublineages of Omicron include BA.1, BA.1.1, BA.2, BA.2.12.1, BA.3, BA.4 and BA.5^38^.

The S glycoprotein homotrimer on the surface of SARS-CoV-2 is critical for viral entry by binding cellular receptor ACE2 and mediating fusion of viral and cell membranes^1,39^. The monomeric S glycoprotein consists of the S1 and S2 subunits. The S1 subunit for receptor binding folds into four major domains including the N-terminal domain (NTD), receptor-binding domain (RBD) and two subdomains (SD1 and SD2), while the S2 for membrane fusion has fusion peptide (FP), two heptad repeats (HR1 and HR2) and other secondary structural elements^40^. SARS-CoV-2 neutralizing antibodies bind the S glycoprotein to block its interaction with the ACE2 receptor or interfere with the pre-fusion to post-fusion conformational transition of the S glycoprotein required for membrane fusion^41,42^. Among the domains and secondary structural elements in the S glycoprotein, RBD is the predominant target of neutralizing antibodies that can be grouped into four classes (class 1 to class 4) based on germline or structural information^43^. By including more antibodies and finer epitope binning, the antibody epitopes on the RBD were further redefined into seven core communities (RBD-1 to RBD-7), which are located on the top receptor-binding motif (RBM) face (RBD-1, RBD-2 and RBD-3), the solvent-exposed outer surface (RBD-4 and RBD-5) and the cryptic inner face (RBD-6 and RBD-7) of the RBD^44^.

Mutations on the RBD play important roles in varied receptor binding and escape of antibody neutralization of SARS-CoV-2 VOCs, thereby affecting viral transmissibility and potency of neutralizing antibodies^29,30,45,46^. Notably, Omicron has close to 20 mutations on the RBD and 10 of them map to the top RBM surface directly interacting with the ACE2 receptor^47,48^. It has been shown that Omicron strikingly reduced or abrogated neutralization titers of sera from vaccinated and convalescent individuals^24,35,49–52^. Most RBD-directed potent antibodies previously identified including those approved for emergency use authorization (EUA) also exhibited markable reduction or complete loss of neutralizing activity against Omicron^24,35,53–56^. For example, a family of class I antibodies using the immunoglobulin heavy chain variable 3-53 or 3-66 gene (IGHV3-53/3-66) strongly bind to the RBM face, and their epitopes are largely within the RBD-2 community and overlap with ACE2-binding site. Majority of them are heavily affected by mutations on the RBM face such as Q493R, Q498R, N501Y and Y505H^43,48,53,56^. Similarly, N440K, G446S and E484A mutations found on the Omicron RBD are involved in reducing the activities of the class 2 and class 3 antibodies targeting RBD-4 and RBD-5 on the outer surface, while S371L, S373P and S375F heavily affect many antibodies in the class 4 targeting RBD-6 and RBD-7 on the inner surface^48,53,56^.

Previously we reported a neutralizing antibody F61 from convalescent patients after prototype SARS-CoV-2 infection, which showed high potency in neutralizing SARS-CoV-2 and Alpha, Beta and Delta variants^57^. In this study, we showed that F61 using the IGHV3-53/3-66 gene exhibited the same high potency in neutralizing Omicron diverse sublineages and protecting mouse model against Delta and Omicron BA.1 variants. Therefore, F61 is an exceptional broadly neutralizing antibody in the family of IGHV3-53/3-66-using antibodies. We also reported another broadly neutralizing antibody D2 that is able to potently neutralize SARS-CoV-2 VOCs except Omicron BA.1.1 and BA.4, although its potency is less than that of F61. More importantly, we showed that F61 and D2 exhibited significant synergy in both in vitro neutralization of all VOCs and in vivo protection against Delta and Omicron BA.1. Cryo-EM structure determination of the spike-Fab complexes revealed the distinct epitopes of F61 and D2 on the RBD and provided structural insights into the broad and potent neutralization by F61 and F61-D2 cocktail against all SARS-CoV-2 VOCs including diverse Omicron sublineages.

## Results

### Biochemical characterization of neutralizing antibodies F61 and D2

We previously reported a phage-displayed antibody library constructed from the PBMCs of SARS-CoV-2 convalescent donors^57^. Both F61 and D2 were isolated by screening this library with wildtype SARS-CoV-2 RBD (WT-RBD). After purifying these two antibodies in the recombinant form of human IgG1, we tested their binding avidity to different fragments of the WT, Delta and Omicron spike glycoproteins using ELISA method. These fragments include WT-S1, WT-NTD, WT-RBD, Delta-S1, Delta-RBD and Omicron-RBD (BA.1). The ELISA results showed that F61 and D2 bound to above fragments but not WT-NTD with EC50 values less than 4.5 ng/mL (Fig. 1A), indicating that they are both RBD specific antibodies.

**Fig. 1.**
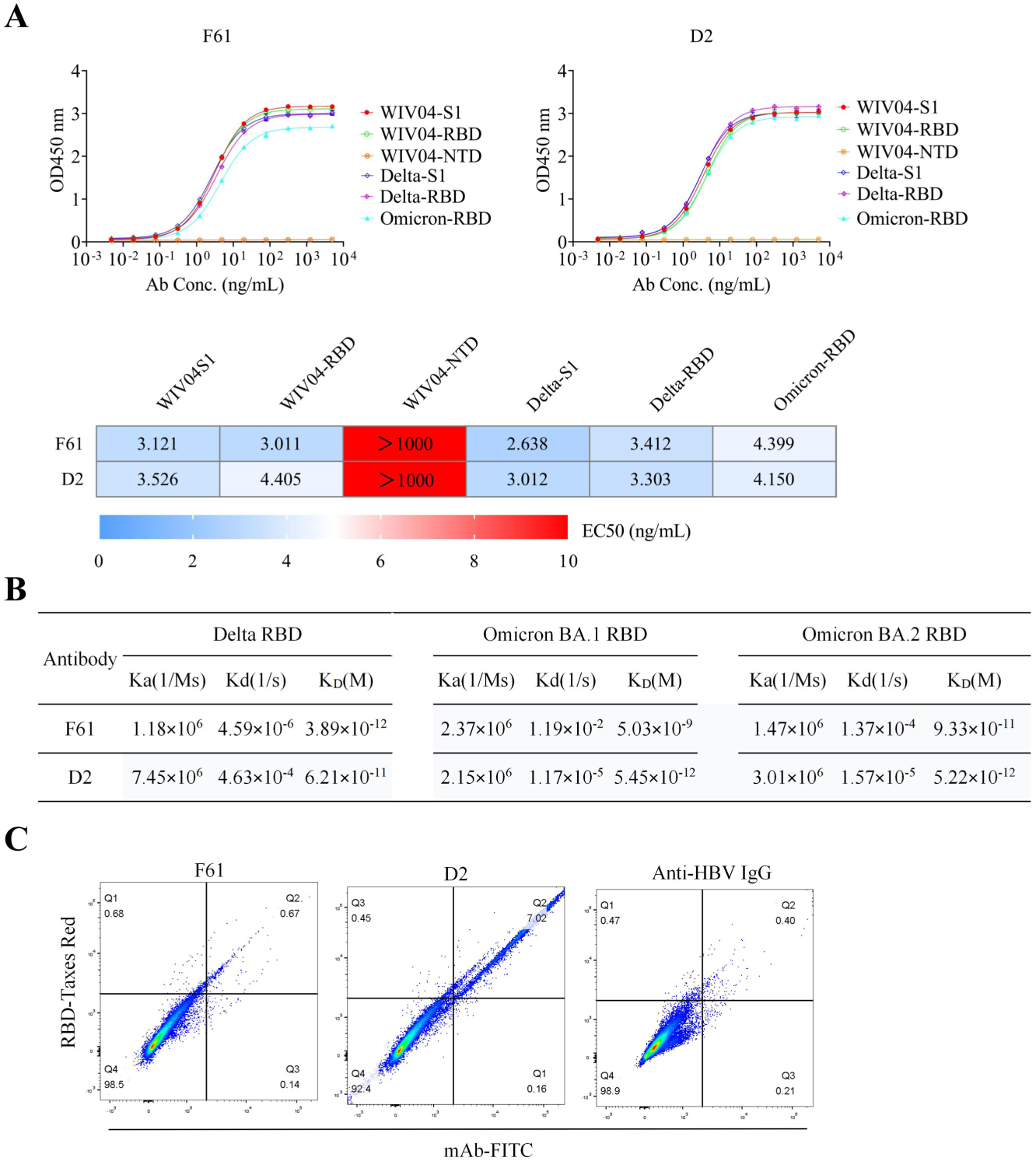
Biochemical characterization of neutralizing antibodies F61 and D2. **A,** Binding profiles of mAbs measured by ELISA. **B,** Binding kinetics of mAbs with Delta-RBD, Omicron-RBD (BA.1 and BA.2) measured by SPR. **C,** Antibody and ACE2 competition for binding to SARS-CoV-2 WT-RBD measured by FACS.

We also measured the binding affinities of F61 and D2 to Delta-RBD and Omicron-RBD (BA.1 and BA.2) using surface plasmon resonance (SPR) method (Fig. 1B and Fig. S1). Both antibodies exhibited high-affinity binding to Delta-RBD, Omicron-RBD-BA.1 and Omicron-RBD-BA.2. The K_D_ values to Delta-RBD, Omicron-RBD-BA.1 and Omicron-RBD-BA.2 by F61 were ∼3.89 pM, ∼5.03 nM and ∼93.3 pM, respectively. We previously reported the K_D_ of ∼3.72 pM to the WT-RBD^57^. Therefore, the binding of F61 to WT-RBD and Delta-RBD are on the same level. Its binding to Omicron-RBD was reduced by ∼1000-fold for BA.1 (K_D_ = ∼5.03 nM) and ∼30-fold for BA.2 (K_D_ = ∼93.3 pM). D2 retained high-affinity binding on pM level to Delta-RBD (K_D_ = ∼62 pM), Omicron-RBD-BA.1 (K_D_ = ∼5.45 pM) and Omicron-RBD-BA.2 (K_D_ = ∼5.22 pM), and the binding to Delta-RBD is slightly weaker than that to Omicron-RBD. Next, we examined the effects of these two antibodies in inhibiting the staining of hACE2-expressing HEK293 cells by WT-RBD-mFC fusion protein using FACS (Fig. 1C). The results showed that F61 not D2 was able to compete with ACE2 receptor in binding WT-RBD, indicating their distinct binding epitopes on the RBD.

### Broadly neutralizing activities of F61 and F61-D2 cocktail against SARS-CoV-2 VOCs in vitro and in vivo

We tested the neutralizing activities of F61 and D2 against pseudoviruses of SARS-CoV-2 WT (D614G) and its VOCs including Alpha (B.1.1.7), Beta (B.1.351), Delta (B.1.617.2 and B.1.617.3) and Omicron (BA.1, BA.1.1, BA.2, BA.3 and BA.4) (Fig. 2A). Corresponding to the high-affinity binding, F61 was highly potent in neutralizing SARS-CoV-2 WT pseudovirus with half-maximal inhibitory concentration (IC50) of 6 ng/mL. All IC50 values against the tested VOCs were less than 20 ng/mL (Fig. 2A). These results showed that F61 exhibited high potency and broad neutralization against Alpha, Beta, Delta and Omicron, even its binding to Omicron-RBD-BA.1 was significantly reduced (∼1000-fold) compared to WT-RBD (Fig. 1B). Similar to F61, D2 was able to potently neutralize SARS-CoV-2 WT pseudovirus with IC50 of 19 ng/mL, and it retained the same level high potencies in neutralizing pseudoviruses of Alpha, Beta, Delta and Omicron including BA.1, BA.2 and BA.3 with IC50 values less than 50 ng/mL (Fig.2A). However, its potency against pseudovirus of Omicron BA.1.1 and BA.4 was reduced with IC50 values increased to 249 and 318 ng/mL, respectively (Fig. 2A). Considering distinct epitopes of F61 and D2 indicated by the competition assay, we also tested the combination of F61 and D2 with a 1:1 molar ratio in pseudovirus neutralization. The results showed that all tested VOCs were well neutralized with IC50 values less than 30 ng/mL by using the F61-D2 cocktail (Fig. 2A).

**Fig. 2.**
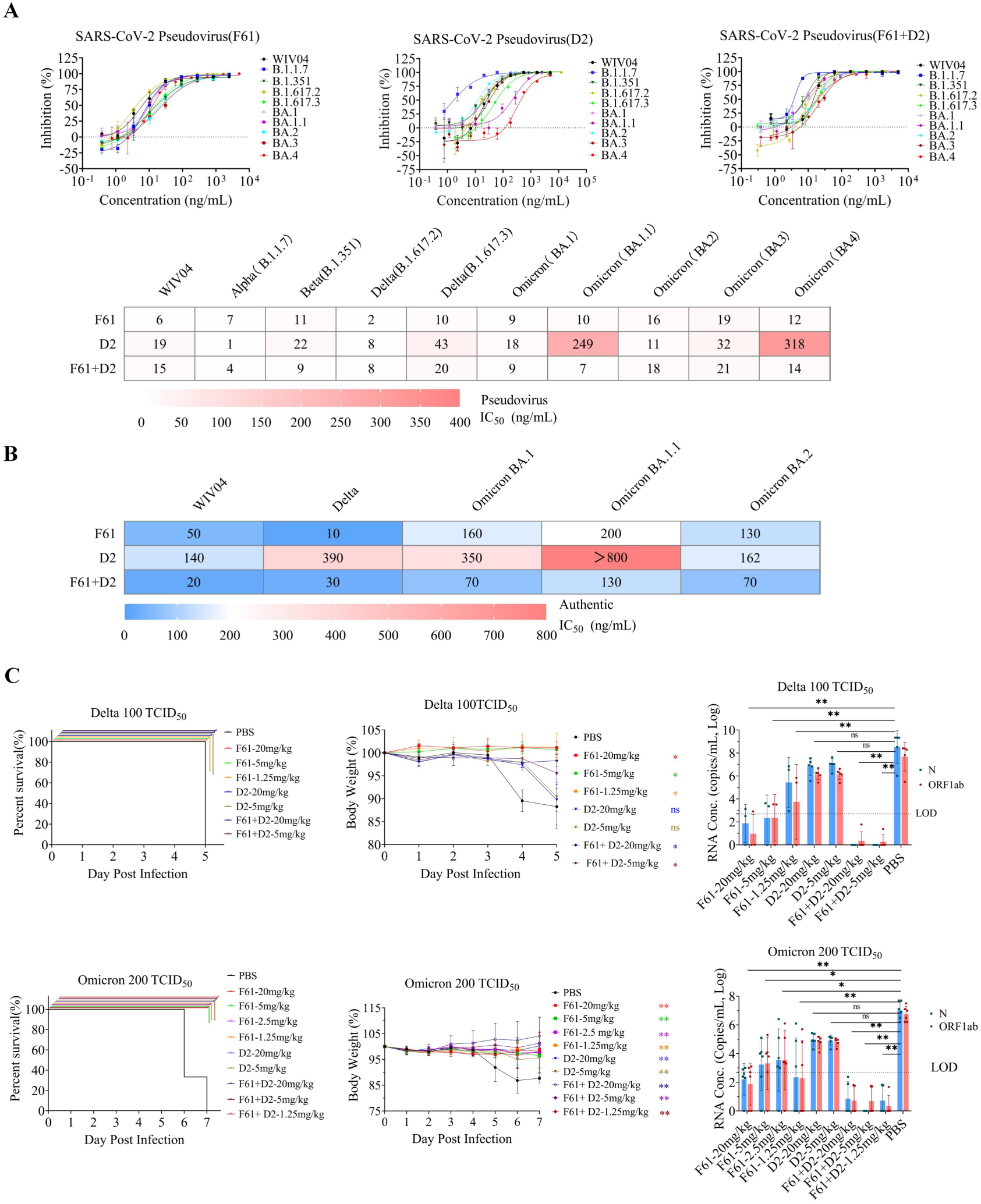
Broadly neutralizing activities of F61 and F61-D2 cocktail against SARS-CoV-2 VOCs in vitro and in vivo. **A**, Neutralization of mAbs to SARS-CoV-2 pseudovirus in HEK293T-hACE2 cells. **B,** Neutralization of mAbs to SARS-CoV-2 authentic virus in Vero E6 cells. **C,** Prophylactic effects of F61, D2 or F61-D2 cocktail against SARS-CoV-2 Omicron BA.1 and Delta variants in K18-hACE2 mice. Body weights change (%), survival curves, and viral RNA loads in the lungs of K18-hACE2 mice treated with difference doses of antibodies (1.25, 2.5, 5, 20 mg/kg) F61, D2, or F61+D2 via intranasal route before infection with 100 TCID_50_/mouse Delta variant (**upper panel**) or 200 TCID_50_/mouse Omicron BA.1 variant (**down panel**). As a negative control, PBS was administered. Body weight curve values represent means ± standard errors of the means (n = 4∼6 mice/group). Significant differences between the antibody treatment group and negative control are shown. All data points for viral load in the lungs are shown, along with the medians. ns, P > 0.05; *, P < 0.05; **, P < 0.0001, as determined by One -way ANONA. Limit of detection (LOD), 500 copies/mL.

Next, we studied the neutralization of authentic SARS-CoV-2, Delta B.1.617.2 and Omicron BA.1, BA1.1 and BA.2 by F61 and D2. F61 had IC50 values of 50, 10, 160, 200 and 130 ng/mL against SARS-CoV-2, Delta, Omicron BA.1, Omicron BA.1.1 and Omicron BA.2, respectively (Fig. 2B). By increasing the IC50 value from 50 ng/mL against SARS-CoV-2 to more than 100 ng/mL against Omicron BA.1, BA.1.1 and BA.2 (Fig. 2B), the effects of Omicron mutations in reducing F61 potency were more obvious in neutralizing authentic viruses than pseudoviruses. The D2 exhibited less potency than F61, with the IC50 values against SARS-CoV-2, Delta B.1.617.2, Omicron BA.1 and BA.2 were 140, 390, 350 and 162 ng/mL, respectively (Fig. 2B). In consistent with reduced activity against Omicron BA.1.1 pseudovirus, its potency against the authentic Omicron BA.1.1 was also significantly impaired with IC50 value of more than 800 ng/mL (Fig. 2B). We further tested the F61-D2 cocktail and the results indicated a synergy between them in the neutralization, especially against Omicron BA.1, BA.1.1 and BA.2. When neutralizing these three Omicron sublineages, F61 and D2 together with a 1:1 molar ratio showed a 1.5 to 6.1-fold improvement in IC50 values over the individual antibodies (Fig. 2B), suggesting an effect that is more than an additive for the F61-D2 cocktail against SARS-CoV-2 and its variants.

Finally, using K18-hACE2 mice as a prophylactic model as previously described^58^, in vivo protective activities of F61, D2 and F61-D2 cocktail were evaluated with the lethal challenge of Delta and Omicron BA.1 viruses, respectively (Fig. 2C). The results showed that, regardless of giving a high dose (20 mg/kg body weight) or low dose (5 mg/kg or 1.25 mg/kg body weight) of F61, D2 or F61-D2 combination, the antibody treatment conferred protection against the lethal challenges with 100 TCID_50_ Delta or 200 TCID_50_ Omicron BA.1 (Fig. 2C, left panel). Most of tested mice in the experimental end-point did not lose their body weight significantly, in particular with Omicron BA.1 challenge (Fig. 2C, middle panel). Moreover, the viral RNAs in the lung of mice of F61 or F61-D2 cocktail groups were significantly reduced or negatively detected compared with PBS groups (10^8^ copies/mL for both Delta or Omicron BA.1), whereas the decrease in viral load in the D2 group was not as significant as in the F61 group (Fig. 2C, right panel). More importantly, the significant synergy between F61 and D2 was also observed in vivo with the lethal challenge of either Delta or Omicron BA.1, even at the minimum administration dose of 1.25 mg/kg body weight, the viral loads in related mice lung were negative or below the minimum detection limit (less than 10^5^ copies/mL) (Fig. 2C, right panel).

### Overall cryo-EM structures of the spike-antibody complexes

To understand structural basis for the binding and neutralization by F61 and D2, we expressed and produced the six proline-stabilized (S6P) WT spike ectodomain with S1/S2 furin-cleavage site mutated to GSAS. The complexes of the WT-spike bound by F61 or D2 Fab were prepared and single particle cryo-EM data were collected, resulting in the binary WT-spike-F61 and WT-spike-D2 structures determined at 3.62 and 3.25 Å, respectively (Fig. S2-S4 and Table S1). We also prepared the Omicron BA.1 spike ectodomain with S6P and GSAS mutations and determined the cryo-EM structure of the ternary Omicron-spike-F61-D2 complex at a resolution of 3.04 Å (Fig. S2-S4 and Table S1). Due to the conformational heterogeneity of the Fab-bound RBDs relative the rest of the spike trimer, only the VH and VL domains of the bound Fab were built in all final models.

It has been found that the apo SARS-CoV-2 spike trimer usually exhibits a mixture of a closed form with all three RBDs in the down position and an open form with one RBD in the up position^40,59,60^. In the binary spike-F61 and spike-D2 complexes, the spike trimer is in the open form with all three RBDs adopting similar upright position with a tilt angle of ∼90 degree (Fig. 3A and 3B). In the spike-F61 complex, each up-RBD was bound by a F61 Fab on the top receptor-binding motif (RBM) surface for ACE2 engagement (Fig. 3A). Among seventeen RBD residues involved in ACE2 binding, ten of them were recognized by F61, resulting in a large overlap between F61 epitope and ACE2 binding site (Fig. 3D). In the spike-D2 complex, each up-RBD was bound by a D2 Fab covering the solvent-exposed outer surface (Fig. 3B), resulting in an epitope on the RBD spatially distinct from ACE2 binding site (Fig. 3D). In the determined Omicron-spike-F61-D2 ternary complex structure, three RBDs are all in the up position, and each up-RBD was bound simultaneously by one F61 Fab and one D2 Fab (Fig. 3C and 3D).

**Fig. 3.**
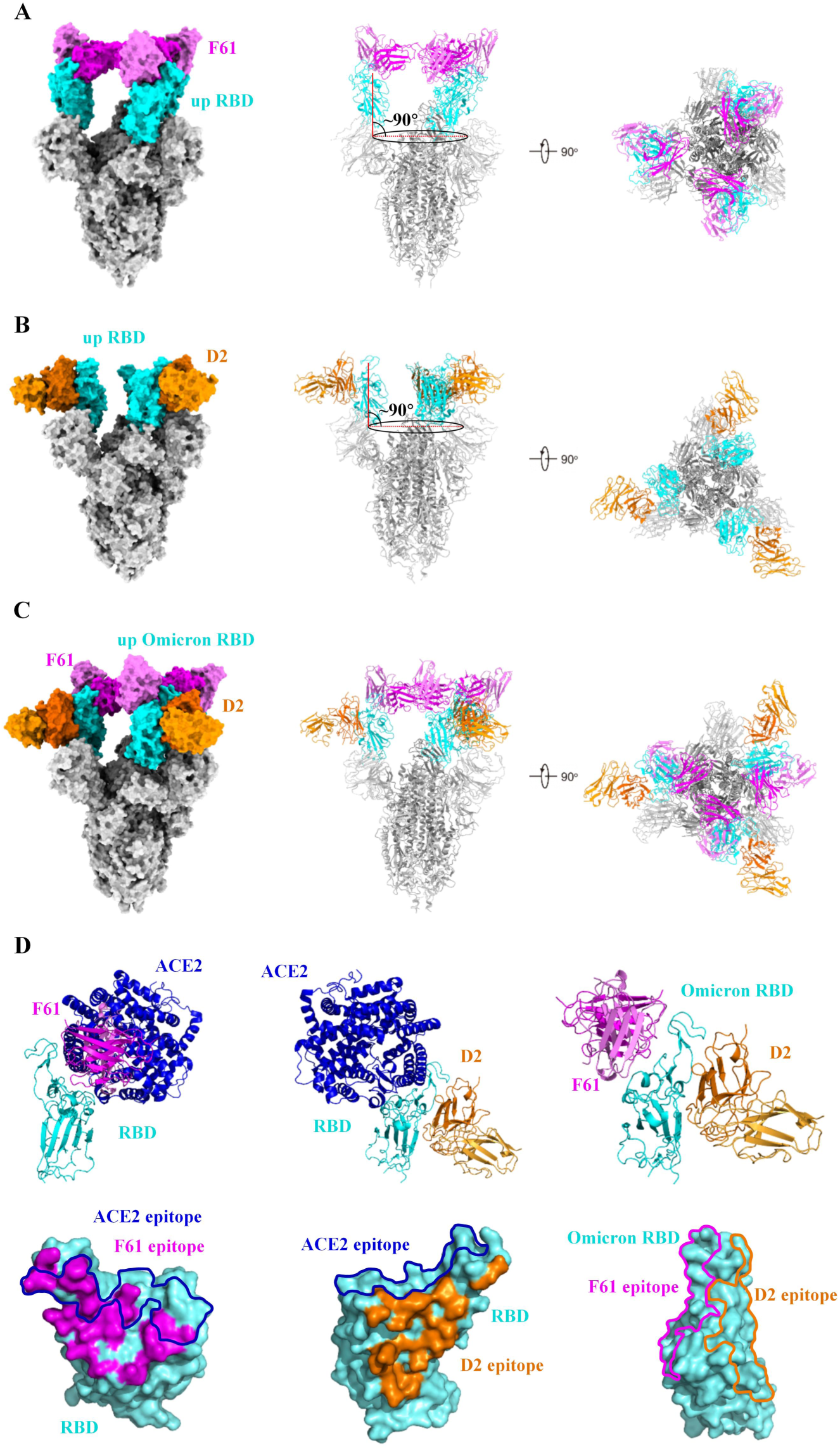
Overall cryo-EM structures of the spike-antibody complexes. **A,** Overall structure of SARS-CoV-2 spike in complex with F61 Fab. The tilt angle of RBD is defined by the angle between the long axis of RBD (red line) and its projection on the horizontal plane (black ellipse). Angle between them is indicated. SARS-CoV-2 RBD is colored in cyan, other domains in grey, heavy chain of F61 in magenta and light chain of F61 in violet. **B,** Overall structure of SARS-CoV-2 spike in complex with D2 Fab. The tilt angle of RBD is defined by the angle between the long axis of RBD (red line) and its projection on the horizontal plane (black ellipse). Angle between them is indicated. SARS-CoV-2 RBD is colored in cyan, other domains in grey, heavy chain of D2 in orange and light chain of D2 in light orange. **C,** Overall structures of SARS-CoV-2 Omicron spike in complex with F61 Fab and D2 Fab. Color schemes are the same as **A** and **B**. **D,** Structural superposition of RBD-fab and RBD-ACE2 (PDB ID: 6M0J) structures and the footprints of F61, D2 and ACE2 on the RBD, and structure of Omicron-RBD-F61-D2 with footprints of F61 and D2 on the RBD. ACE2 is colored in blue. The footprints of F61, D2 and ACE2 are represented as magenta, orange and blue, respectively. Other color schemes are the same as **A** and **B**.

### Structural basis for the potent and broad neutralization

We performed focused 3D classification and local refinement of the Fab-RBD region, resulting in improved densities for building the WT-RBD/Fab and Omicron-RBD-BA.1/Fab interfaces (Fig. S4). We utilized the Omicron-RBD-BA.1/Fab interfaces for the following structural description and analysis, due to their better densities compared to WT-RBD/Fab interfaces. F61 using IGHV3-66 gene is a member of the class 1 antibody, and it binds to an epitope on the RBM face that can be grouped into the RBD-2a community^43,44^ (Fig. 4A). The interaction buried a total of 1145 Å^2^ surface area from F61 and 1075 Å^2^ from RBD. All six CDRs of F61 are involved in RBD binding (Fig. 4A), and the heavy chain is more dominant than the light chain by contributing 19 residues among all 26 antibody residues for binding (Table S2). The F61 epitope consisting of 25 RBD residues does not include Alpha mutation and includes Beta K417N and Delta T478K substitutions. Omicron has more mutations in the F61 epitope, including K417N, S477N, T478K, Q493R and Y505H for BA.1 and BA.1.1. F61 epitope includes additional mutation sites found in other Omicron sublineages, which are D405N and R408S on BA.2, D405N on BA.3 and D405N, R408S and F486V on BA.4 and BA.5 (Fig. 4A and 4C). At the Omicron-RBD-BA.1/F61 interface, N417, N477, K478, R493 and H505 have extensive interactions with F61 residues E26, I28, Y33, Y99, D101 and F102 from the heavy chain, and N31 and D51 from the light chain (Fig. 4C). Therefore, these Omicron mutations work in concert to alter specific interactions at the interface and significantly reduced the binding of F61 to Omicron-RBD-BA.1 (K_D_ = ∼5.03 nM) compared to WT-RBD (K_D_ = ∼3.89 pM) (Fig. 1B). Compared to Omicron-RBD-BA.1, the binding of Omicron-RBD-BA.2 to F61 was restored to some extent with the K_D_ value of ∼93.3 pM (Fig. 1B), indicating that the additional D405N and R408S mutations on Omicron BA.2 would play roles in enhancing the binding by F61. To be note, even for Omicron-RBD-BA.1 whose binding by F61 was the most significantly reduced (∼1000-fold), the affinity between them is still in low nM range (∼5.03 nM). We conclude that the tight binding and direct ACE2 competition would allow for the high potency and broad neutralization of F61 to be retained against SARS-CoV-2 VOCs including diverse Omicron sublineages.

**Fig. 4.**
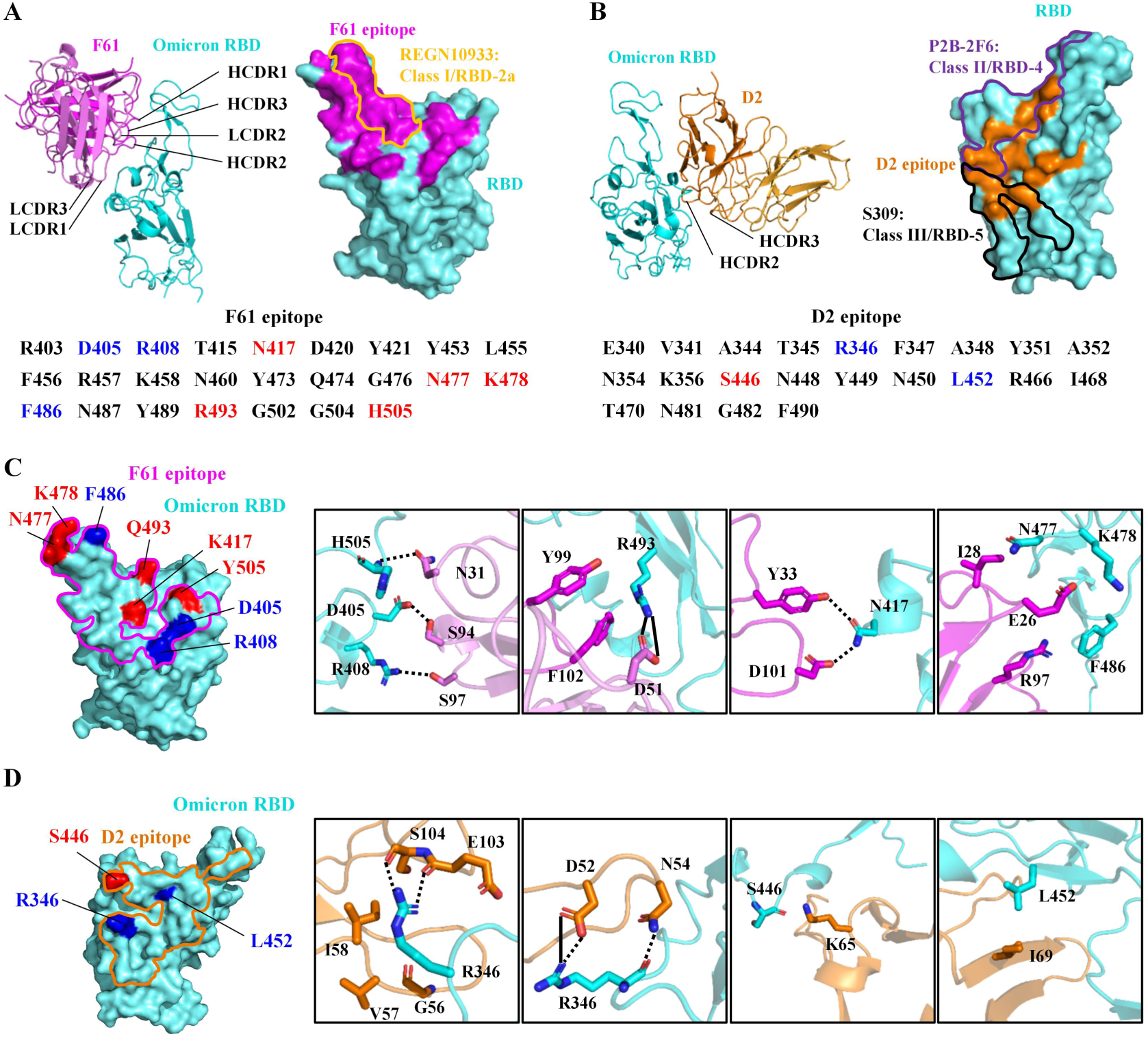
Structural basis for the potent and broad neutralization. **A,** Overall structure of Omicron-RBD-BA.1 bound with F61 and CDRs involved in binding are labeled. On the RBD, the footprints of F61 is represented by magenta surface and the footprint of REGN10933 (PDB ID: 6XDG) is circled by yellow line. Omicron-RBD-BA.1 residues recognized by F61 are listed and mutated N417, N477, K478, R493 and H505 in BA.1 are colored red. Additional mutations sites in BA.2 (D405 and R408), BA.3 (D405), BA.4 (D405, R408 and F486) and BA.5 (D405, R408 and F486) are colored blue. **B,** Overall structure of Omicron-RBD-BA.1 bound with D2 and CDRs involved in binding are labeled. On the RBD, the footprint of D2 is represented by orange surface and the footprints of P2B-2F6 (PDB ID: 7BWJ) and S309 (PDB ID: 6WPS) are circled by purple and black lines, respectively. Omicron-RBD-BA.1 residues recognized by D2 are listed and mutated S446 in BA.1 is colored red. Additional mutation site in BA.1.1 (R346) and only L452 mutation site in BA.4 and BA.5 are colored blue. **C,** The detailed interactions between residues mutated in above Omicron sublineages and F61. F61 footprint is circled by a magenta line. Interacting residues of Omicron-RBD are shown as cyan sticks and F61 are shown as magenta sticks. Hydrogen bonds and salt bridges are represented by dashed lines and solid lines, respectively. **D,** The detailed interactions between residues mutated in above Omicron sublineages and D2. D2 footprint is circled by an orange line. Interacting residues of Omicron-RBD are shown as cyan sticks and D2 are shown as orange sticks. Hydrogen bonds and salt bridges are represented by dashed lines and solid lines, respectively.

Structure determination showed that the epitope D2 using IGHV3-9 gene is on the outer surface of the RBD, locating between previously defined RBD-4 and RBD-5 communities and having overlap with both of them^43,44^ (Fig. 4B). Upon binding, the VH domain of D2 contacts the up-RBD by aligning in parallel with the outer surface, whereas the VL domain does not have contact with the RBD (Fig. 4B). The interaction between D2 VH domain and up-RBD buried a total of 953 Å^2^ surface area from the VH domain and 944 Å^2^ from the RBD. At the interface, 11 residues from the D2 heavy chain HCDR2 and HCDR3 interact with 13 Omicron-RBD residues (Fig. 4B and Table S2). Among close to 20 mutations on the Omicron-RBD, the D2 epitope include only G446S mutation found on BA.1 but not on BA.2 (Fig. 4D). The interaction around S446 at the Omicron-RBD-BA.1/D2 interface is between heavy chain K65 and the main-chain oxygen atom of S446 (Fig. 4D), which could explain that the binding of D2 to Omicron-RBD BA.1 or BA.2 was retained on pM level and D2 could still potently neutralize Omicron BA.1 and BA.2 (Fig. 1B, Fig. 2A and 2B). D2 epitope includes G446S and R346K mutation sites on BA.1.1 and only L452R on BA.4 and BA.5 (Fig. 4B & 4D). R346 has extensive interactions with D2 residues D52, N54, G56, V57, I58, E103 and S104, including hydrogen bonds of R346 with D52, N54, E103 and S104 and salt bridge of R346 with D52 (Fig. 4D). R346K substitution is expected to abolish some hydrogen bonds and to interfere with interactions around the 346 between Omicron BA.1.1 RBD and D2, resulting in reduced neutralization against Omicron BA.1.1 compared to BA.1, BA.2 and BA.3 (Fig. 2A and 2B). L452 has hydrophobic interaction with D2 heavy chain I69 (Fig. 4D). Similar to R346K mutation, the L452R mutation found in Omicron BA.4 would also interfere local interactions and result in reduced neutralization against BA.4 compared to BA.1, BA.2 and BA.3 (Fig. 2A and 2B).

## Discussion

A large number of potent neutralizing antibodies against SARS-CoV-2 have been reported since the beginning of the COVID-19 pandemic^4,6–9,13,14,20^. However, the emergence of SARS-CoV-2 VOCs harboring mutations on the spike glycoprotein has led to great concerns over resistance to neutralizing antibodies and failure of vaccines^23,24,30,35,49–56,61^. In fact, recent studies have found that most of previously identified neutralizing antibodies have shown markable reduction or complete loss of activities against Omicron^24,35,53–56^. Here we comprehensively characterized a highly potent and broadly neutralizing antibody F61 and its cocktail with another antibody D2 against SARS-CoV-2 VOCs including Omicron.

The IGHV3-53/3-66-using antibodies are frequently elicited in most people after SARS-CoV-2 infection or vaccination and many of them exhibit high potency by strongly binding to the RBM and directly competing with ACE2 receptor^62^. However, most of them are heavily affected by VOCs, especially Omicron carrying multiple mutations on the RBM^24,53–56,63^. Previously we showed that F61 was highly potent against SARS-CoV-2 WT and Alpha, Beta and Delta variants in pseudovirus inhibition^57^. Here we showed that F61 was still highly potent against Omicron BA.1, BA.1.1, BA.2 BA.3 and BA.4 pseudoviruses with IC50 values below 20 ng/mL. It was able to neutralize cell infection of authentic Omicron BA.1, BA.1.1 and BA.2 with IC50 values below 200 ng/mL, respectively. Structure of Omicron-RBD-BA.1/F61 interface indicated that despite close to twenty mutations in Omicron-RBD-BA.1, F61 could still interact with these residues, such as N417, N477, K478, R493 and H505. Furthermore, F61 also formed salt bridges with R493 and hydrogen bonds with N417 and H505. Given that neutralization titers of many monoclonal antibodies and sera from vaccinated and convalescent individuals are significantly reduced because of K417N, Q493R and Y505H mutations, robust binding with these mutation sites by F61 may explain why it stands out from other antibodies and shows excellent neutralization against all tested variants, including Omicron BA.1, BA.1.1, BA.2, BA.3 and BA.4 sublineages. These results collectively showed that F61 is an exceptional IGHV3-53/3-66-using antibody exhibiting potent and broadly neutralizing activity. CAB-A17 is another recently reported IGHV3-53/3-66-using antibody with broad neutralizing activity^64^. CAB-A17 and F61 exhibited similar high potency against Omicron pseudovirus (CAB-A17 IC50: ∼15 ng/mL; F61 IC50: 10-20 ng/mL)^64^. Structure and sequence comparisons showed that the Omicron-RBD epitope residues are nearly the identical, and antibody residues involved in hydrogen-bonding interaction are also highly conserved between CAB-A17 and F61, such as heavy chain E26, Y33, G54, S56 and R97^64^. The study of CAB-A17 also found that only four somatic hypermutations G26E, T28I, S53P and Y58F were able to confer breadth to CAB-A17 to against Omicron^64^, and E26, I28, P53 and F58 are also conserved in F61. To be note, F61 and CAB-A17 are two significant examples of VOC-neutralizing antibodies isolated from convalescent patients before circulation of VOCs, indicating that broadly neutralizing antibodies are within the repertoire after prototype SARS-CoV-2 infection and such memory B cells could be recruited upon re-infection or vaccination.

Here we also reported another broadly neutralizing antibody D2, although its potency is weaker than that of F61. Structural elucidation of the RBD/D2 interfaces confirmed that the epitope of D2 is relatively conserved and almost unchanged in tested VOCs, and thus it is unaffected by most VOCs. It only includes G446S mutation in Omicron BA.1, BA.2 and BA.3 and our structure shows that D2 remains interaction with Omicron BA.1 S446. However, RBD residue R346 is within the epitope and has extensive interactions with D2, which may dramatically reduce its neutralization activity against Omicron BA.1.1. These structural observations fit well with functional data showing that D2 broadly bound and neutralized Omicron BA.1, BA.2 and BA.3 and its potency was significantly reduced against Omicron BA.1.1 carrying the R346K mutation. Although RBD-directed SARS-CoV-2 neutralizing antibodies have been extensively studied and summarized such as the class 1-4 antibody and RBD1-7 communities^43,44^, the overall binding mode and epitope of D2 are still out of ordinary. Its epitope is between RBD-4 and RBD-5 and overlaps with both of them. Therefore, unlike many class 2 antibodies having RBD-4 epitope, D2 does not compete with ACE2 and its epitope does not include E484K/A mutation that reduces or abolishes the neutralizing activity of many antibodies in the RBD-4 community^53,54,56^. At the same time, unlike many class 3 antibodies having RBD-5 epitope, D2 does not bind to the N343-linked glycans centered in RBD-5 epitope, which is highly conserved among SARS-CoV-2, SARS-CoV-1 and many bat and pangolin betacoronaviruses^14,44,65^. Structural comparison showed that D2 is similar to a recently reported antibody COVOX-58 in overall binding mode and epitope on the RBD^61^ (Fig. S5). These two antibodies use the same IGHV3-9 gene. Although the HCDR3 of COVOX-58 is longer than that of D2, their epitopes on the RBD are very similar due to the dominant role of the conserved HCDR2 in RBD binding by F61 and COVOX-58^61^ (Fig. S5).

We also proved that F61 and D2 exhibited significant synergy both in vitro and in vivo. Animal experiments showed that F61-D2 cocktail could provide protection against Delta and Omicron BA.1. Even when mice were administrated at the minimum dose of 1.25 mg/kg body weight, the viral loads in their lung were still negative or below the minimum detection limit. These in vivo experiments implicate that F61-D2 cocktail could be a promising therapeutic combination for combating SARS-CoV-2 variants of concern, including Omicron. We aligned RBD-F61 or RBD-D2 binary complexes onto the spike trimers in the closed form or in the open form with one or two RBDs adopting the up conformation. The results showed that F61 and D2 epitopes are only fully exposed in the up-RBD (Fig. S6), indicating that the down to up conformational change is a prerequisite for the binding of F61 and D2. After binding to the RBM face, one important mechanism of F61 neutralization is to directly block spike-ACE2 interaction. The major neutralizing mechanism of D2 would not be ACE2 competition because its epitope does not overlap with ACE2-binding site. The destabilization of spike trimer by representative class 3 antibody S309 was suggested to be its mechanism of action^48^, and D2 may utilize similar working mechanism. Due to the distinct epitopes, when F61 and D2 are used as a cocktail, one antibody binding to the RBD may help to induce and fix the RBD in the up conformation for efficient binding of the other antibody. In this way, the F61-D2 cocktail would be more efficient than individual F61 in fully binding the spike trimer and blocking all ACE2-binding sites, as shown in the Omicron-RBD-F61-D2 ternary complex structure (Fig. 3C).

## Figures and Figure legends

**Fig. S1.**
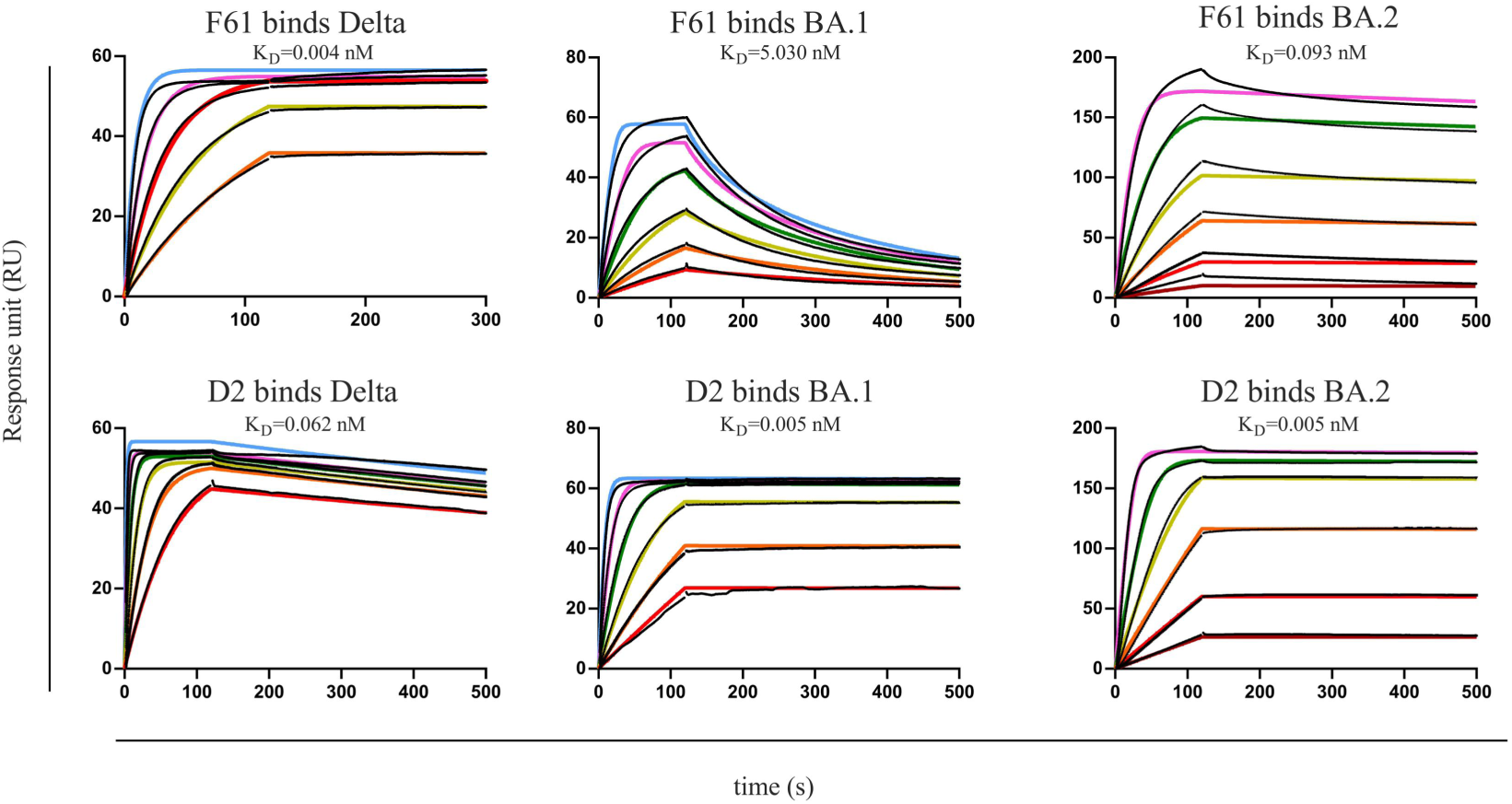
The binding curves between mAbs and SARS-CoV-2 Delta or Omicron RBD measured by SPR. Black lines were the original curves, while colored lines were the fitted curves.

**Fig. S2.**
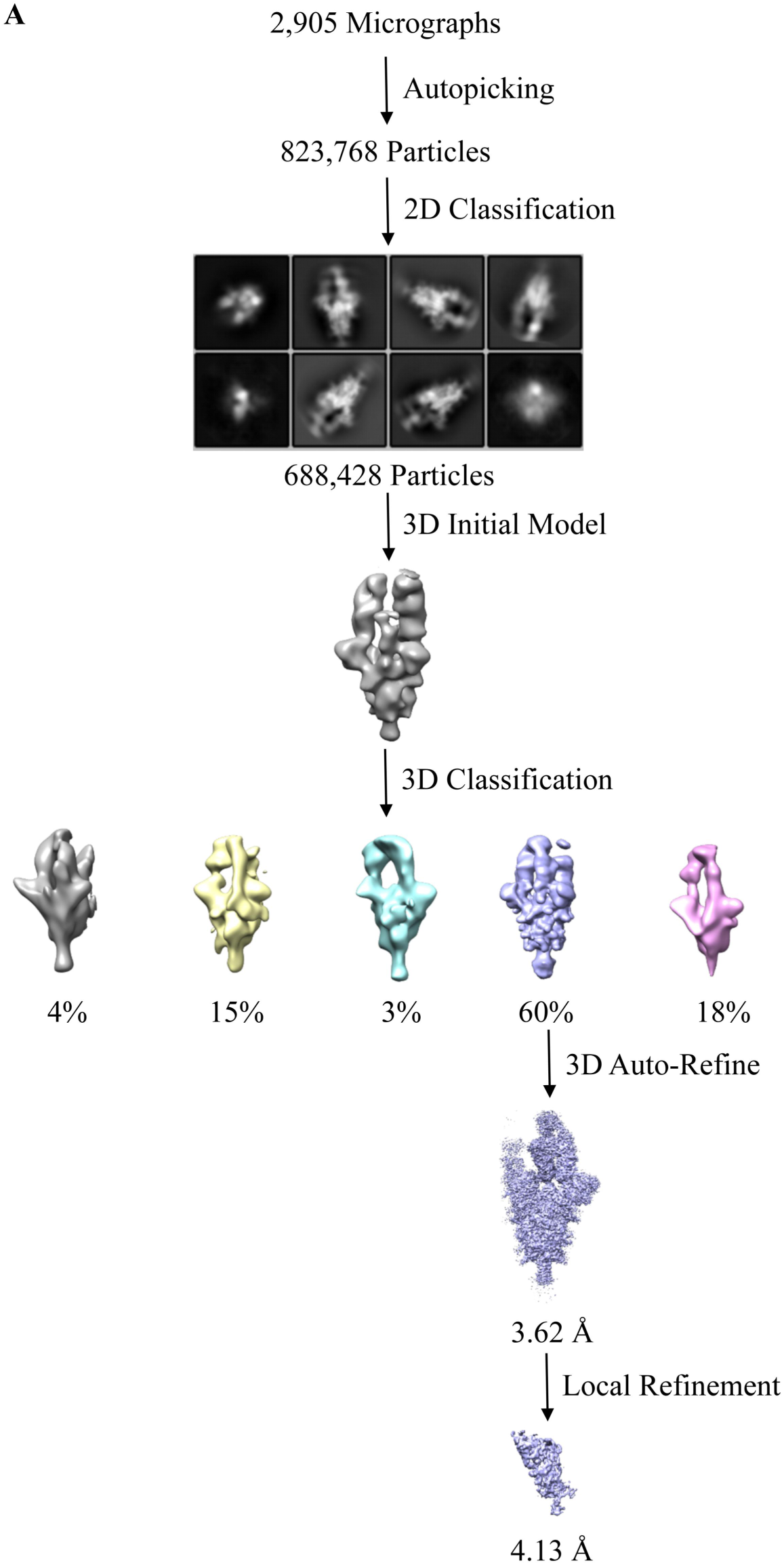
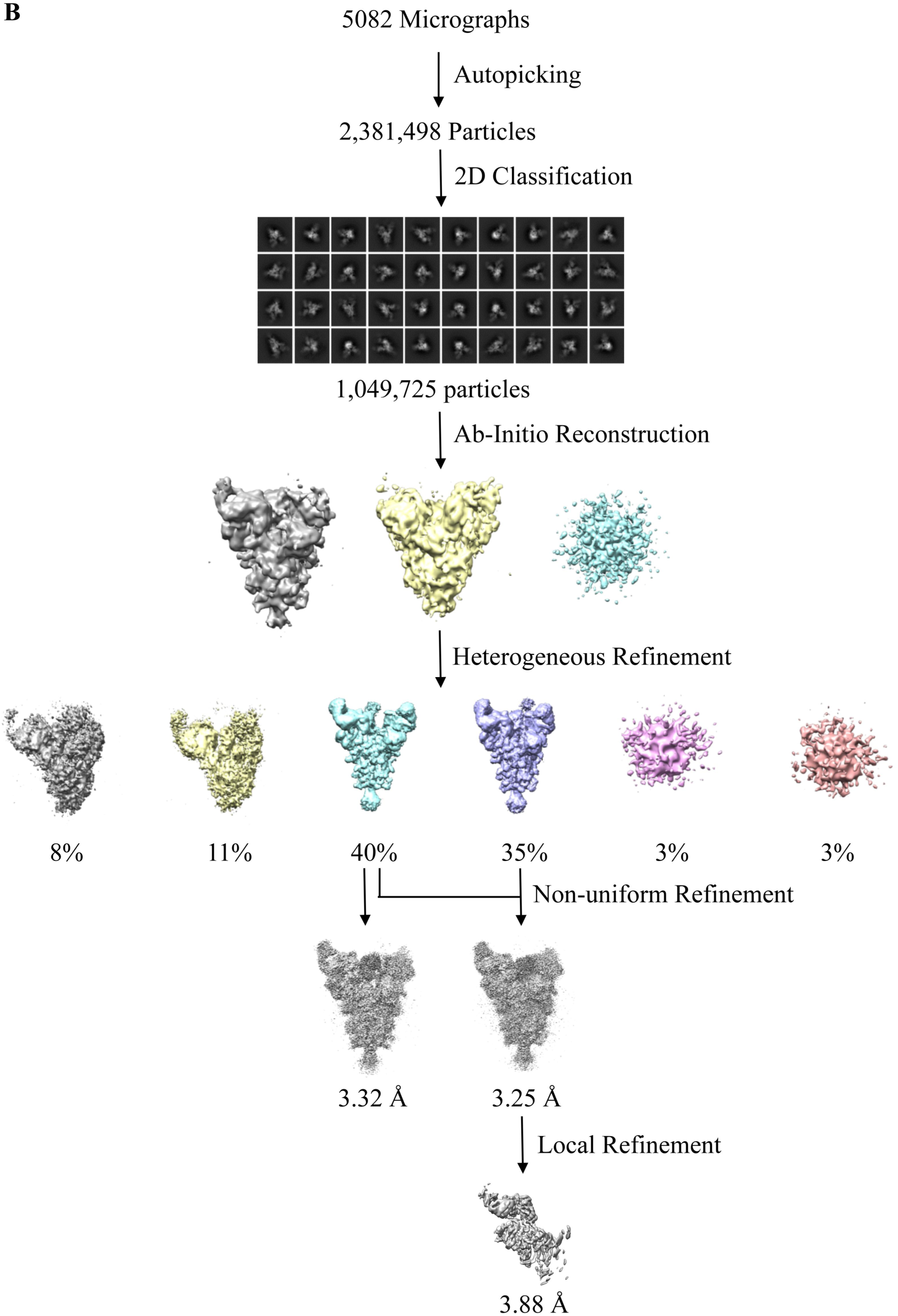
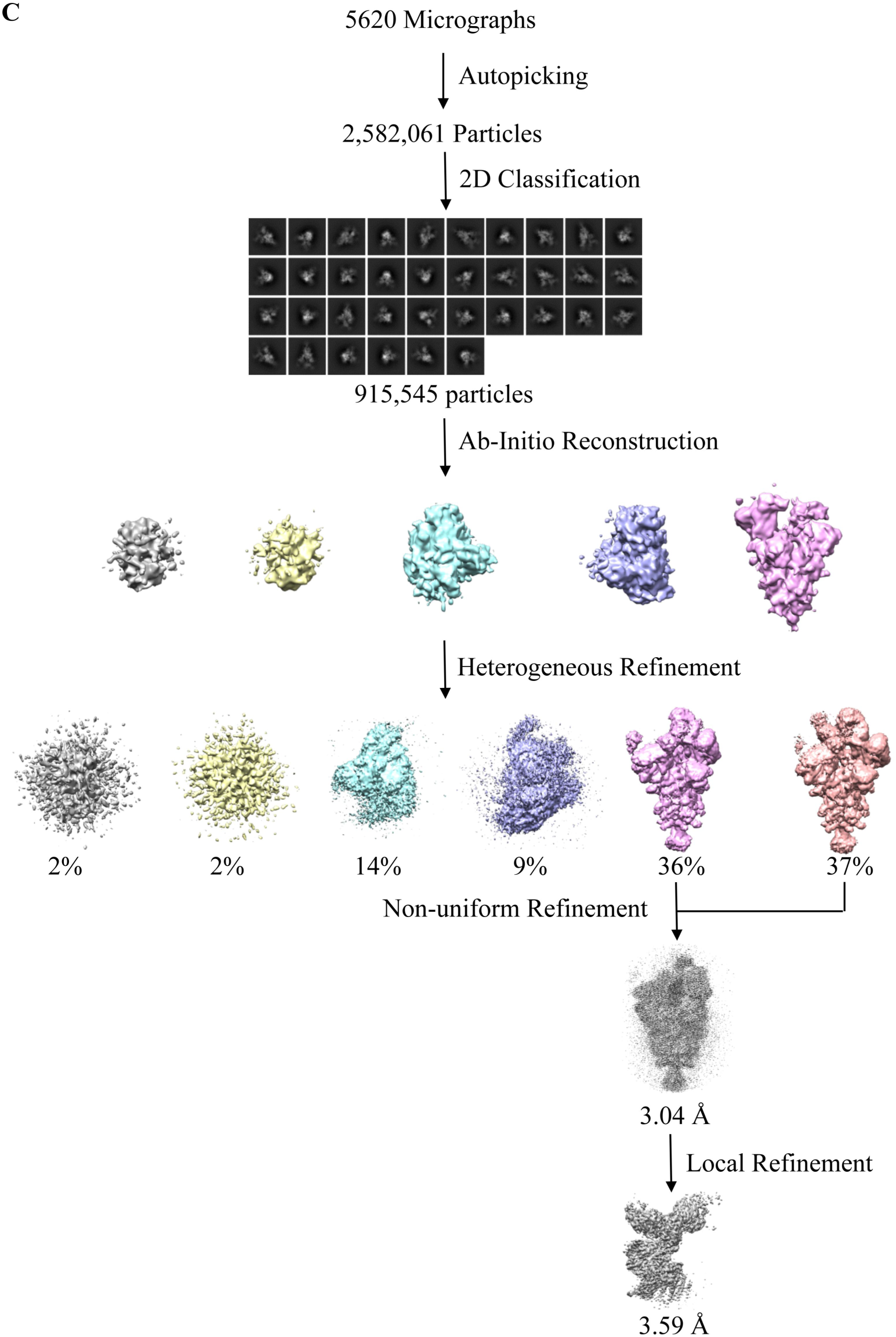
Cryo-EM data processing workflow. **A,** Processing workflow of the Spike-F61 binary complex Cryo-EM data. **B,** Processing workflow of the Spike-D2 binary complex Cryo-EM data. **C,** Processing workflow of the Omicron-Spike-F61-D2 ternary complex Cryo-EM data.

**Fig. S3.**
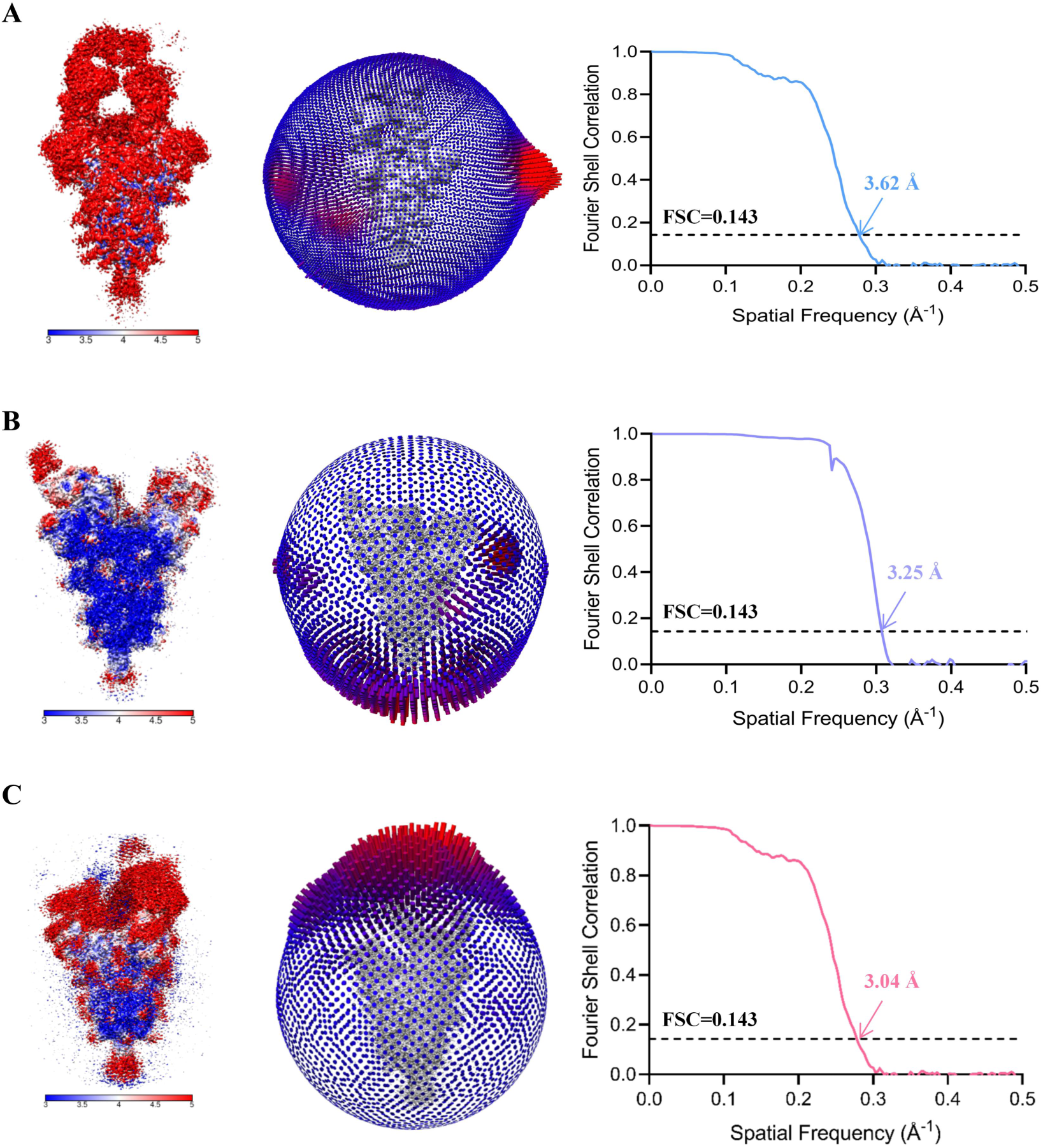
Cryo-EM structure validations. **A-C,** Local resolution map (left panel), particle orientation distribution (middle panel) and Gold-standard Fourier Shell Correlation (FSC) curves of the final density maps (right panel) of the Spike-F61 complex (**A**), the Spike-D2 complex (**B**) and the Omicron-Spike-F61-D2 complex (**C**). The final resolution of the Spike-F61 complex is 3.62 Å, the final resolution of the Spike-D2 complex is 3.25 Å, and the final resolution of the Omicron-Spike-F61-D2 complex is 3.04 Å.

**Fig. S4.**
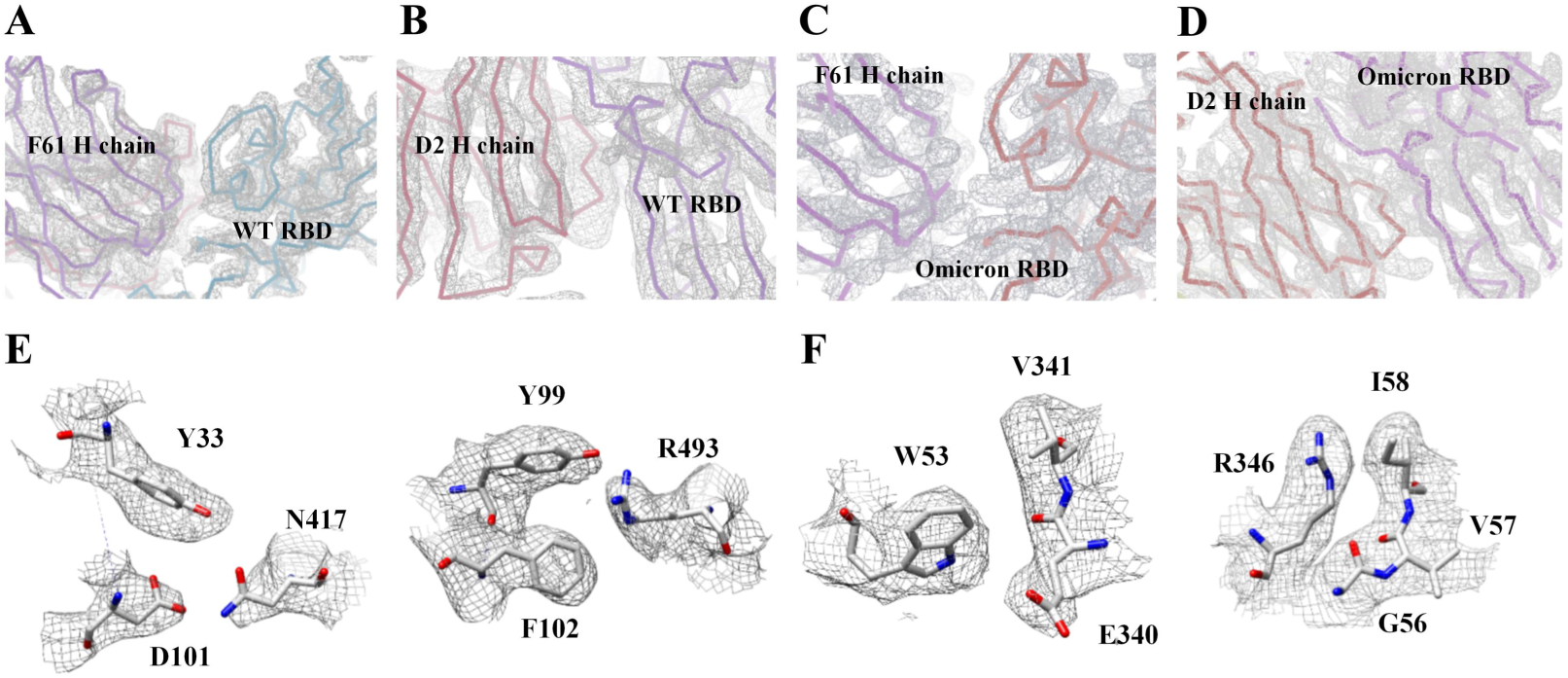
The representative density maps of residues of the Spike-F61, Spike-D2 and Omicron-Spike-F61-D2 complexes. **A,** The representative density maps of the WT-RBD-F61 interface. The map is contoured at 2 RMS to show the density. **B,** The representative density maps of the WT-RBD-D2 interface. The map is contoured at 7.2 RMS to show the density. **C,** The representative density maps of the Omicron-RBD-F61 interface. The map is contoured at 5.6 RMS to show the density. **D,** The representative density maps of the Omicron-RBD-D2 interface. The map is contoured at 5.6 RMS to show the density. **E,** The representative density maps of the Omicron-RBD-F61 interacting residues. The map is contoured at 2.2 RMS to show the density. **F,** The representative density maps of the Omicron-RBD-D2 interacting residues. The map is contoured at 2.2 RMS to show the density.

**Fig. S5.**
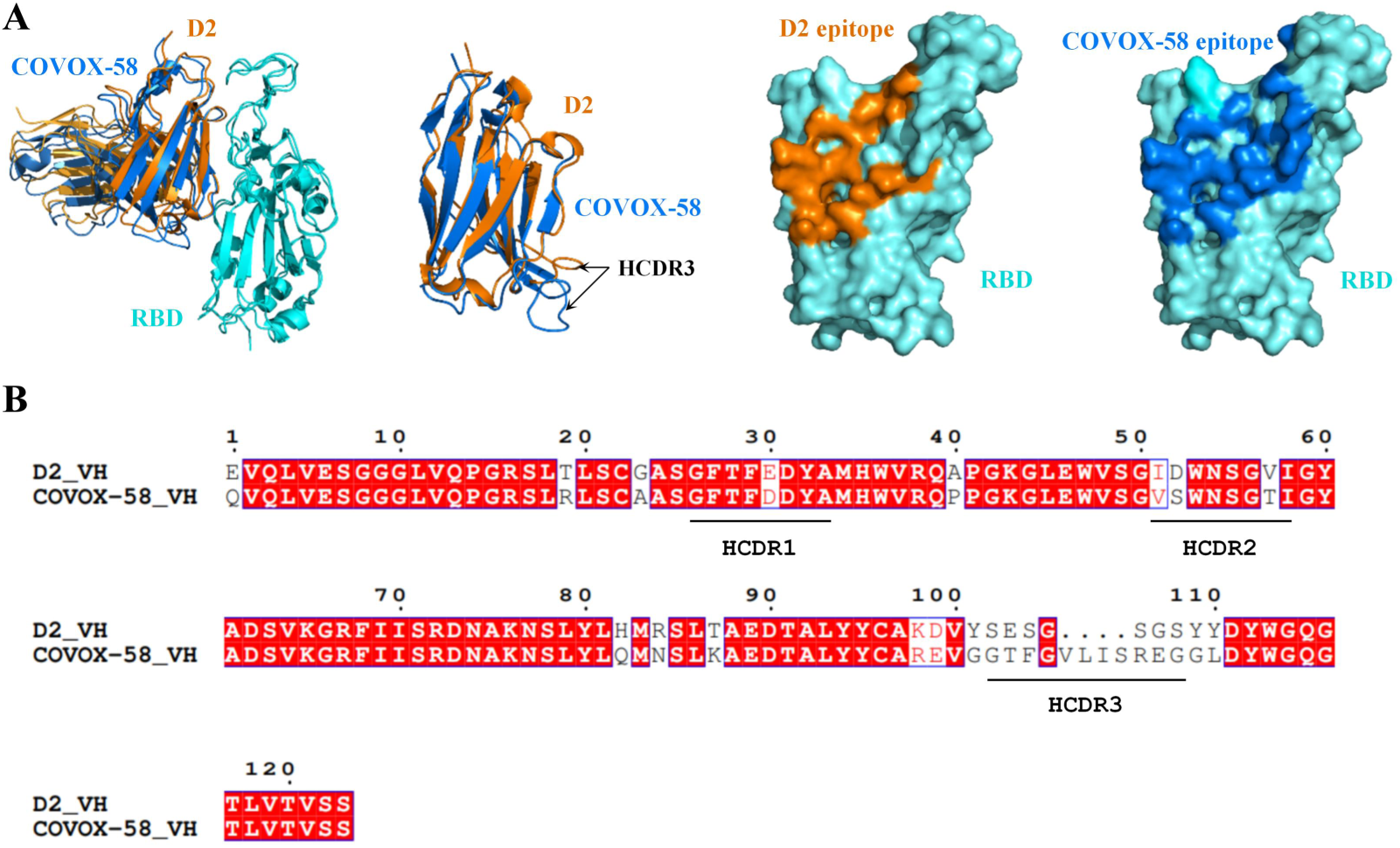
The comparison of RBD-D2 and RBD-COVOX-58 structures and VH sequences. **A,** The structural comparison of the RBD-D2 and RBD-COVOX-58 (PDB ID: 7QNY) complexes. These two structures are superimposed to show a similar binding mode and nearly identical epitope. The structure and epitope of COVOX-58 is shown in marine. HCDR3s of D2 and COVOX-58 are indicated with arrows. **B,** Sequence alignment of VH of D2 and COVOX-58.

**Fig. S6.**
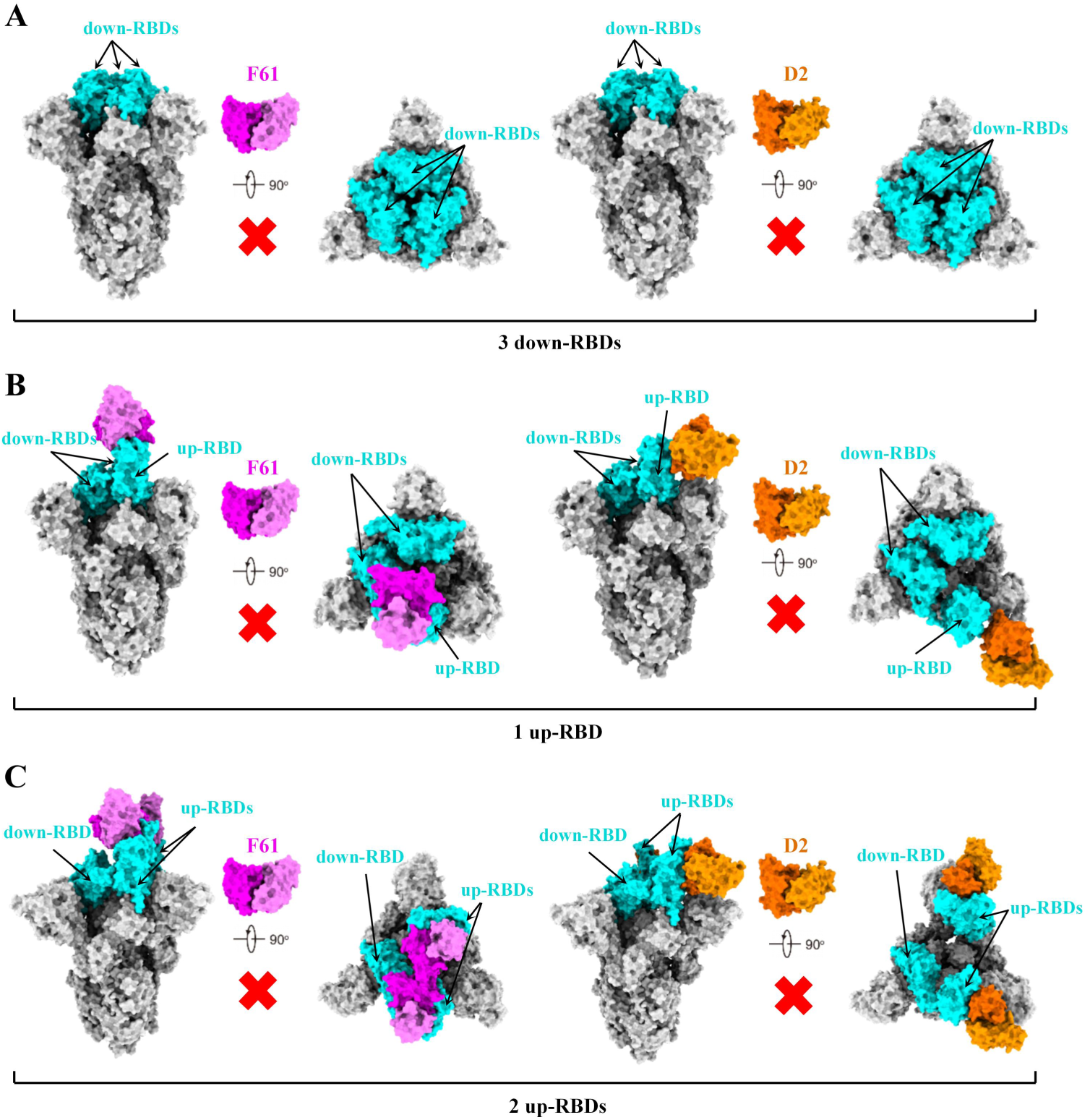
Alignment of the RBD-F61 or RBD-D2 structures onto the spike trimers in the closed form or in the open form with one or two RBDs adopting the up conformation. **A-C,** Align RBD-F61 or RBD-D2 complexes to spike trimer with three RBDs in the down conformation (**A**, PDB ID: 6VXX), spike trimer with one RBD in the up conformation (**B**, PDB ID: 6VSB) and spike trimer with two RBDs in the up conformation (**C**, PDB ID: 7A93).

**Table S1.**
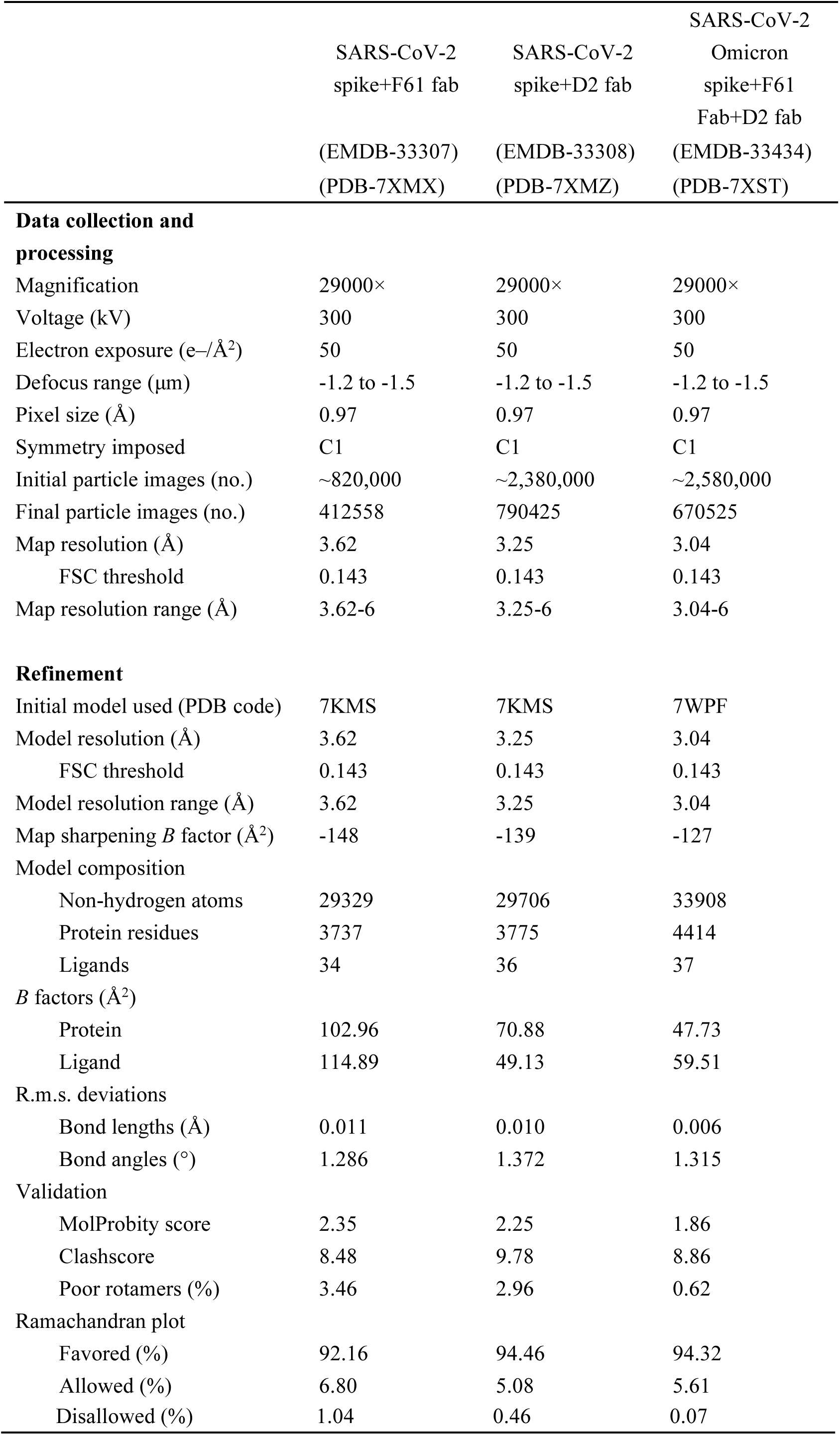
Cryo-EM data collection, refinement and validation statistics.

**Table S2.**
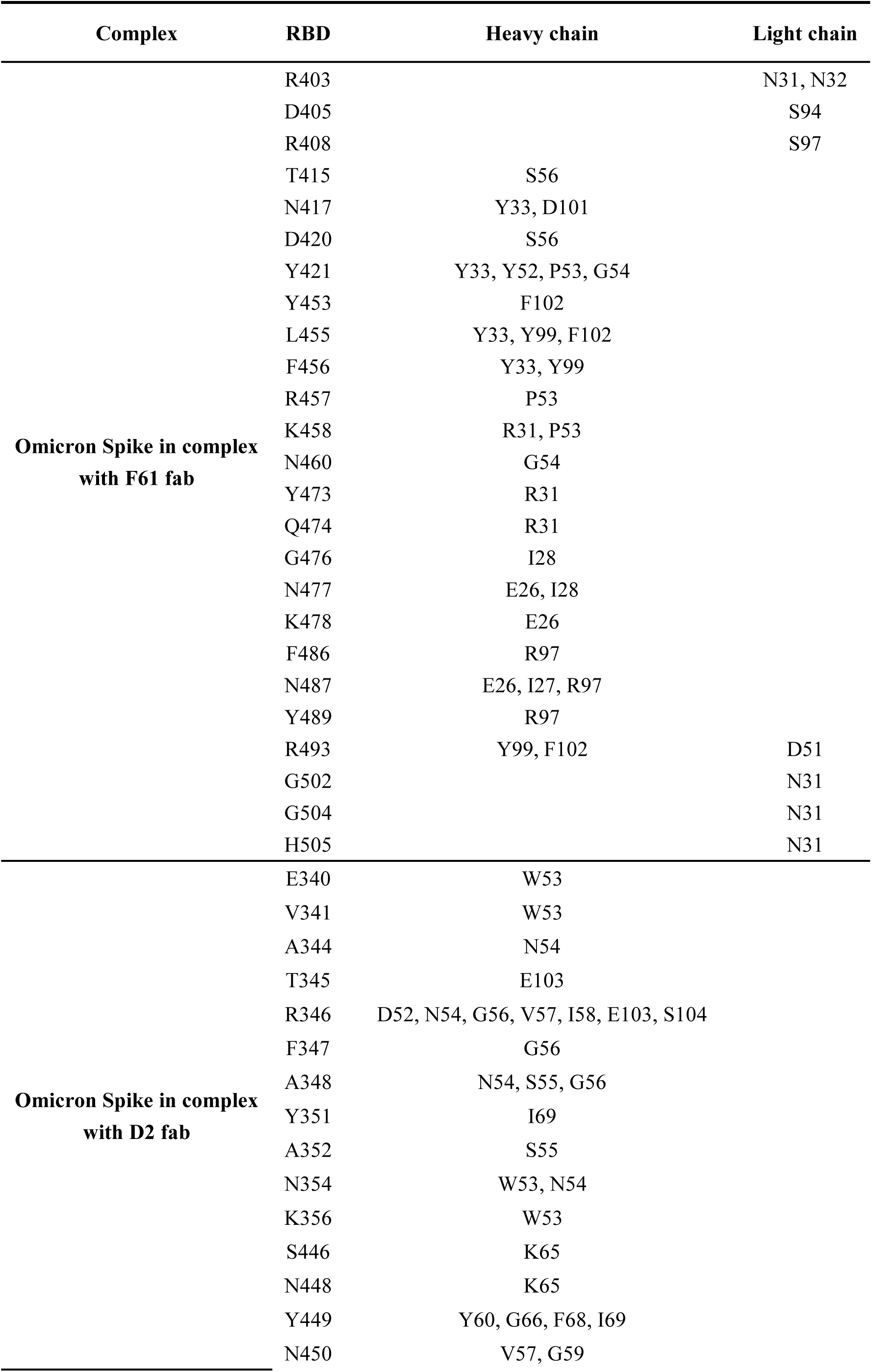
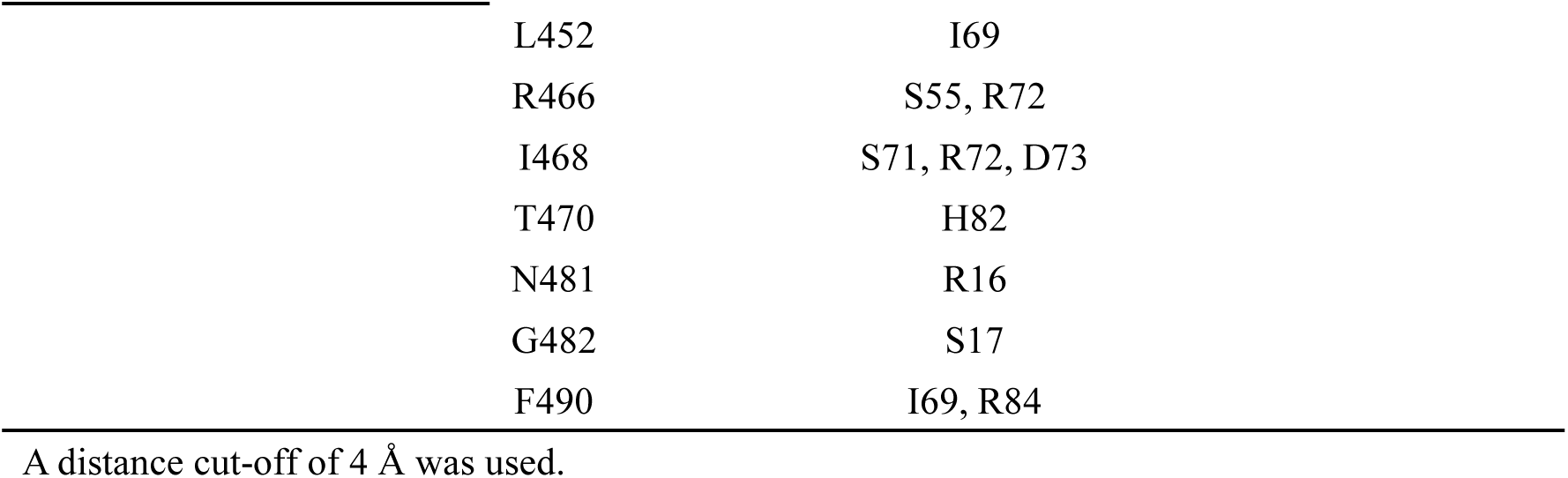
The interacting residues between Fabs and SARS-CoV-2 Omicron Spike.

## Materials and Methods

### Cells, Viruses and Proteins

Cell lines (HEK293T and Vero E6 cells) were initially acquired from the American Type Culture Collection (ATCC; USA). HEK293T-hACE2-cells were generated via the overexpression of the human ACE2 receptor in HEK293T cells and were used in the neutralization assays of pseudoviruses. Vero E6 cells were used in the neutralization assay of authentic viruses. SARS-CoV-2 WT and variants pseudoviruses were purchased from Beijing Tiantan Pharmaceutical Biotechnology Development Co., Ltd. All SARS-CoV-2 authentic virus were isolated from nasopharyngeal and oropharyngeal samples from patients with COVID-19 and deposited by Wuhan Institute of Biological Products Co., Ltd. Recombinant SARS-CoV-2 proteins, including WT-S1/RBD/NTD (Sino, 40591-V49H\40591-V08H\40592-V02H1), Delta (B.1.617.2) S1/RBD with a his tag (Sino, 40591-V08H23\40592-V08H90), Omicron (BA.1) RBD with a his tag (Sino, 40592-V08H121), Omicron (BA.2) RBD with a his tag (ACRO Biosystems, SPD-C522g) were used in the context of phage-display library panning, binding ELISA or SPR.

### Binding ELISA

ELISA plates were coated with SARS-CoV-2 protein including WT-S1, WT-NTD, WT-RBD, Delta-S1, Delta-RBD and Omicron-RBD (Sino Biological, China) at 4 °C overnight. Following washing with PBST, serial dilutions of testing antibodies start at 5 μg/mL were added to each well and incubated at 37 °C for 30 min. After washing with PBST, horseradish peroxidase (HRP)-conjugated anti-human IgG Fc specific antibody (Sigma, USA) was added at the dilution of 1:2,000 and incubated at 37 °C for 30 min. The absorbance was detected at 450 nm. The data were analyzed using GraphPad Prism 8.0.

### RBD-ACE2 Binding Inhibition Assayed by FACS

The block assay was assessed by FACS. HEK293T cells were transiently transfected with the ACE2 expression plasmid for 24 h. The mouse-Fc tag Fusion protein of SARS-CoV-2 RBD (RBD-mFC) (Jiangsu East-MabBiomedical Technology, China) at a concentration of 2 μg/mL was mixed with the mAbs or isotype IgG hepatitis b virus (HBV) at a molar ratio of 1:10 and incubated at 4 °C for 1 h. Then mixtures were added to 2.5 × 10^5^ HEK293T cells expressing ACE2 and incubated at 4 °C for another hour. Then cells were stained with anti-mouse IgG Taxes red conjugated antibody and anti-human IgG FITC-conjugated antibody (Sigma, USA) for another 30 min then analyzed by FACS Aria II (BD, USA). All of these data were analyzed using Flow Jo.

### Antibody Binding Kinetics Measured by SPR

The binding kinetics of mAbs to SARS-CoV-2 Delta-RBD or Omicron-RBD monomer were analyzed using SPR (Biacore 8K; GE Healthcare). Specifically, recombinant protein A (Sino Biological) was immobilized to a CM5 sensor chip. The mAbs (2 μg/mL) were captured by recombinant protein A, and then serial dilutions of SARS-CoV-2 Delta/omicron-RBD with highest concentration of 100 nM to 50 nM were running at a flow rate of 30 μL/min in PBST buffer (1×PBS and 0.05% [vol/vol] Tween-20). The resulting data were fitted to a 1:1 binding model using the Biacore 8K Evaluation software (GE Healthcare). The equilibrium dissociation constants (binding affinity, K_D_) for each antibody were calculated using Biacore 8000 Evaluation Software.

### Virus Neutralization Assay

The neutralization of authentic SARS-CoV-2 reference strain (GenBank ID: MN996528.1) and variants including Delta B.1.617.2 and Omicron BA.1, BA1.1 and BA.2 were measured by the microneutralization test in the bio-safety Level 3 (BSL-3) laboratory. The assay was performed as described by Manenti et al. with a few modifications^66^. Briefly, two-fold serially diluted antibodies (50 µL) in minimal essential medium (Gibco, Thermo Fisher Scientific, Waltham, MA, USA) supplemented with two percent fetal bovine sera (Gibco, Thermo Fisher Scientific, Waltham, MA, USA) were prepared (four replicates per dilution). In the next step, 50 µL of virus suspension of 100 tissue culture infective dose of previously titrated virus stock was added to each well of a 96 well plate (Greiner bio-one GmbH, Frickenhausen, Germany) and incubated at 37 °C for one hour. 100 µL of Vero E6 cells (1 × 10^5^ cells/mL) was then added to the 96 well plates and incubated at 37 °C with 5% CO_2_. After incubation for 72 h, cytopathic effect (CPE) was observed under a light microscope (Nikon, ×100, Tokyo, Japan). The number of positive holes in each row was counted and the neutralizing titer was calculated using the Reed-Muench method.

Neutralization activity of monoclonal antibodies against SARS-CoV-2 pseudoviruses were assayed as previously described^67^. 50 μL serial dilutions of monoclonal antibodies (mAbs) were added into 96-well plates. After that, 50 μL SARS-CoV-2 WT or variants pseudoviruses were incubated with mAbs at 37 °C for 1 h. HEK293-hACE2 cells (2.5 × 10^4^ cells/100μL per well) were then added into the mixture and incubated at 37 °C in a humidified atmosphere with 5% CO_2_ for 23 h to 25 h. Then the luciferase activity was measured after cell lysis. The percent of neutralization was determined by comparing with the virus control. The half-maximal inhibitory concentrations (IC50) were determined using 4-parameterlogistic regression (GraphPad Prism version 8).

### In vivo Protection Activity Evaluation

To evaluate the prophylactic effects of monoclonal antibodies, SARS-CoV-2 Delta and Omicron strains were used in challenge experiments. Male K18-hACE2 mice (6–8 weeks old, purchased from GemPharmatech Co., Ltd. Company.) were randomly distributed into groups (n = 3–6 mice per group). 2 h after administration of monoclonal antibodies, mice were anesthetized with isoflurane and then administered 50 µL SARS-CoV-2 via intranasal route in a challenge dose of 100 or 200 TCID_50_/mouse respectively for Delta and Omicron strains. 50 µL of the antibodies at different concentrations (1.25, 2.5, 5 and 20 mg/kg F61, D2 or F61-D2 (ratio of 1:1)) or vehicle (PBS) was administered to each mouse via intranasal route at 2 h before challenge. Mice were monitored every day for body weight changes and clinical signs of disease until all the mice in the control group died. Mice that lost greater than or equal to 25% of their initial body weight were humanely euthanized. At the end point of the experiment, all remaining animals in the monoclonal antibody-administered group received an overdose of isoflurane and were humanely euthanized. Lungs were collected from each mouse postmortem. Tissues were stored at −80 °C until further analysis.

Tissue homogenates were generated using the TissueLyzer II (Qiagen, Gaithersburg, MD, USA). Briefly, 1000 µL PBS was added to each sample (lungs, 0.01–0.04 g) along with Tungsten carbide 3 mm beads (Qiagen). Samples were homogenized at a speed of 10 Hz for 10 min and then centrifuged at 15,000 × *g* for 10 min. Supernatant was collected, aliquoted, and stored at −80 °C until further analysis.

Total RNA was extracted from tissues homogenates of lungs using an RNA/DNA Purification Kit (Magnetic Bead) (cat no. DA0623; Daan Gene Co., Ltd., China), and RT-qPCR was performed using a Detection Kit for 2019-nCoV (PCR-Fluorescence) (Fast) (cat no. DA0992; Daan Gene Co., Ltd.) following the manufacturer’s instructions. Samples were processed in duplicate using the following cycling protocol: 50 °C for 2 min, 95 °C for 2 min, followed by 42 cycles at 95 °C for 5 s and 60 °C for 10 s. Viral RNA concentrations (copies/mL) in the lungs of mice were determined using RNA standards for SARS-CoV-2 (Bdsbiotech Co., Ltd., Guangzhou, China). The RT-qPCR results were read according to the Daan Kit criteria, the negative results in this manuscript description mean no signal detected (0 copy) or the CT values of both *N* and *ORF1ab* genes were over 38, corresponding viral RNA copies were under the limitation of the detection (LOD, 10^2.7^ copies/mL). All statistical analysis was performed using GraphPad Prism 8. All statistical tests were described in the relevant figure legends.

### Ethics statement

This study was approved by the Experimental Animal Welfare and Ethical Review Board of Wuhan Institute of Biological Products Co., Ltd (protocol WIBP-AⅡ442021005). The experiments conducted in strict accordance with the recommendations in the Guide for the Care and Use of Laboratory Animals established by the People’s Republic of China.

### Expression and purification of SARS-CoV-2 Spike ectodomain and SARS-CoV-2 Omicron Spike ectodomain

The cDNA encoding SARS-CoV-2 WT Spike (GenBank ID: QHD43416.1) was synthesized. Its codons were optimized for insect cell expression and there were six sites mutated to proline. These substitutions occurred at F817, A892, A899, A942, K986 and V987. Furthermore, ‘GSAS’ substitutions were introduced to residues 682-685, the S1/S2 furin cleavage site. The SARS-CoV-2 Spike ectodomain (1-1208) with a C-terminal Strep tag for purification and a foldon tag for trimerization was inserted into the pFastBac-Dual vector (Invitrogen) and was expressed using Bac-to-Bac baculovirus system (Invitrogen). The constructed recombinant plasmid was transformed into bacterial DH10Bac competent cells, then the extracted bacmid was transfected into Sf9 insect cells using Cellfectin II Reagent (Invitrogen). 7 days later, the baculoviruses were harvested. The low-titre viruses were then used for amplification to generate high-titre baculoviruses, which were used to infect Hi5 insect cells at a density of 2 × 10^6^ cells per ml for protein expression. 60 hours after infection, the supernatant of cell medium containing SARS-CoV-2 Spike was collected and concentrated with buffer changed into Tris buffer (50 mM Tris, pH 8.0, 150 mM NaCl). The SARS-CoV-2 spike ectodomain was purified by Strep-Tactin beads (IBA) and eluted with 10 mM Desthiobiotin in Tris buffer. Then the interest protein was purified by gel-filtration chromatography using a Superose 6 gel filtration column (GE Healthcare) pre-equilibrated with HBS buffer (10 mM HEPES, pH 7.2, 150 mM NaCl). Fractions containing the SARS-CoV-2 Spike ectodomain were collected and concentrated for subsequent electron microscopy analysis.

The cDNA encoding SARS-CoV-2 Omicron Spike was synthesized (GenBank ID: ULC25168.1) and cloned into the pCAG vector. There were six sites mutated to proline and these substitutions occurred at F817, A892, A899, A942, K986 and V987. Furthermore, ‘GSAS’ substitutions were introduced to the S1/S2 furin cleavage site. The SARS-CoV-2 Omicron Spike ectodomain (1-1213) with a C-terminal Strep tag for purification and a foldon tag for trimerization was expressed in FreeStyle 293-F cells (Invitrogen). The plasmid was transiently transfected at a density of 2 × 10^6^ cell per ml using polyethyleneimine (PEI) (Sigma) with a mass ratio of 1:4, and the supernatant was collected 4 days later. The supernatant was concentrated with buffer changed into Tris buffer (50 mM Tris, pH 8.0, 150 mM NaCl). The SARS-CoV-2 Omicron Spike ectodomain was purified by Strep-Tactin beads (IBA) and eluted with 10 mM Desthiobiotin in Tris buffer. Then the protein was purified by gel-filtration chromatography using a Superose 6 gel filtration column (GE Healthcare) pre-equilibrated with HBS buffer (10 mM HEPES, pH 7.2, 150 mM NaCl). Fractions containing the SARS-CoV-2 Omicron Spike ectodomain were collected and concentrated for subsequent electron microscopy analysis.

### Preparation of Fab fragments

F61 and D2 Fab fragments were prepared by digesting F61 and D2 IgG with papain (Sigma), respectively. And then Protein A beads (GenScript) were used to separate Fab fragments, following by gel-filtration chromatography using a Superdex 200 column (GE Healthcare) pre-equilibrated with HBS buffer.

### Cryo-electron microscopy sample preparation, data collection and processing

The purified SARS-CoV-2 Spike ectodomain was mixed with F61 and D2 Fab with a molar ratio of 1:3, respectively. The purified SARS-CoV-2 Omicron Spike ectodomain was mixed with F61 and D2 Fab with a molar ratio of 1:3:3. The final concentrations of the three mixtures were 0.82, 1.37 and 0.91 mg/mL in HBS buffer, respectively. Then, Spike trimer - Fab complexes (4 μl) were applied to the pre-glow-discharged holey carbon grids (Quantifoil grid, Cu 300 mesh, R1.2/1.3). The grids were then blotted for 2 seconds with filter paper in 100% relative humidity and 8 °C and plunged into the liquid ethane to freeze samples using FEI Vitrobot system (FEI).

Cryo-EM data were collected using FEI Titan Krios (Thermo Fisher Scientific) electron microscope operating at 300 kV with a Gatan K3 Summit direct electron detector (Gatan Inc.) at Tsinghua University. 2905 movies were collected for Spike-F61 complex, 5082 movies were collected for Spike-D2 complex and 5620 movies were collected for Omicron-Spike-F61-D2 complex using the SerialEM software^68^. These data were collected at a magnification of 29,000 with a pixel size of 0.97 Å and at a defocus range between 1.2-1.5 μm. Each movie had a total accumulate exposure of 50 e^-^/Å^2^ fractionated in 32 frames of 66 ms exposure.

MotionCor2 v.1.2.6^69^ was used for beam-induced motion correction of whole frames in each movie, and GCTF v.1.18^70^ was used to estimate the parameters of contrast transfer function (CTF) for each micrograph. Particles were automatically picked using Gautomatch (http://www.mrc-lmb.cam.ac.uk/kzhang/). And ∼820,000 particles for Spike-F61 complex, ∼2,380,000 particles for Spike-D2 complex and ∼2,580,000 particles for Omicron-Spike-F61-D2 complex were extracted using RELION 3.0.8^71^, which were used for subsequent 2D classification. Spike-F61 complex used RELION 3.0.8^71^ for subsequent data processing. Spike-D2 complex and Omicron-Spike-F61-D2 complex used cryoSPARC^72,73^ for subsequent data processing. After one or two rounds of 2D classification, the preferable classes were selected and these selected particles were used to create 3D initial model and perform 3D classification. Finally, a total of 412,558 particles for Spike-F61 complex, 790,425 particles for Spike-D2 complex and 670,525 particles for Omicron-Spike-F61-D2 complex were applied to 3D refinement to generate density map and post-processing was performed. Based on the gold-standard Fourier shell correlation (FSC) cutoff of 0.143 criterion, the resolutions were 3.62 Å for Spike-F61 complex, 3.25 Å for Spike-D2 complex and 3.04 Å for Omicron-Spike-F61-D2 complex. Local refinement was then performed to further improve the density of the interaction interface of the spike and the Fabs. Local resolution variations were estimated using ResMap 1.1.4^74^. Data collection and processing statistics of Spike-F61 complex, Spike-D2 complex and Omicron-Spike-F61-D2 complex were listed in Table S1.

### Model building and refinement

The structure of the SARS-CoV-2 Spike in complex with ACE2 with 3-up RBDs (PDB: 7KMS) was used to generate the initial model of Spike for Spike-F61 and Spike-D2 structures, the structure of the SARS-CoV-2 Omicron Spike in complex with JMB2002 with 3-up RBDs (PDB: 7WPF) was used to generate the initial model of Omicron Spike for Omicron-Spike-F61-D2 structure, and the initial model of Fabs were predicted using AlphaFold2^75^. These atomic models were fit into the final density maps using UCSF Chimera v.1.16^76^. Coot v.0.9.2^77^ was subsequently used for manual adjustment and correction according to the protein sequences, map densities, Ramachandran plot, rotamers and bond geometry restraints. The Real Space Refinement of PHENIX v.1.18.2^78^ was also used to refine these structures. The quality of the final models was evaluated by PHENIX v.1.18.2^78^. The validation statistics of these structural models were listed in Table 1. Figures were generated using PyMOL 2.0.7^79^, UCSF Chimera v.1.16^76^, UCSF ChimeraX v.1.13^80^.

## Acknowledgments

We thank Dr. Yuhe Yang for generously providing the plasmid and protein of SARS-CoV-2 Omicron Spike. We thank the Tsinghua University Branch of China National Center for Protein Sciences (Beijing) for technical support (Cryo-EM, Protein Preparation and Characterization and Biocomputing). This work was supported by funds from the National Key Plan for Scientific Research and Development of China (2021YFC2300104, 2017YFA0205100 and 2021YFC2600200), the National Natural Science Foundation of China (32171202) and Tsinghua University Spring Breeze Fund (2020Z99CFY031).

## Author contributions

X.W., X.Y., M.L. and K.D. conceived and designed the study. X.L. carried out protein purification, cryo-EM sample preparation, data collection and processing, model building and refinement with the help of L.Z. and J.Y.. X.L. analyzed the structural data and made related figures. S.S., Y.Q., F.G., J.L. Z.J. and T.H. performed the biochemical experiments and provided original data. Q.Y., L.S. and W.W. analyzed data. Y.P., Z.W., X.S. and J.L. contributed reagents, materials and analysis tools. X.L., D.L., S.W. and L.Z. analyzed and discussed the data. X.L., Q.Y., L.S. and W.W. wrote the manuscript, and X.W., X.Y., M.L. and K.D. edited the manuscript.

## Data availability statements

The coordinates of SARS-CoV-2 Spike-F61, SARS-CoV-2 Spike-D2 and SARS-CoV-2 Omicron Spike-F61-D2 have been deposited in the Protein Data Bank (PDB) with the accession numbers 7XMX, 7XMZ and 7XST, respectively; their corresponding maps have been deposited in the Electron Microscopy Data Bank (EMDB) with the accession numbers EMD-33307, EMD-33308 and EMD-33434, respectively.

